# The plot thickens: haploid and triploid-like thalli, hybridization, and biased mating type ratios in *Letharia*

**DOI:** 10.1101/2020.12.18.423428

**Authors:** S. Lorena Ament-Velásquez, Veera Tuovinen, Linnea Bergström, Toby Spribille, Dan Vanderpool, Juri Nascimbene, Yoshikazu Yamamoto, Göran Thor, Hanna Johannesson

## Abstract

The study of the reproductive biology of lichen fungal symbionts has been traditionally challenging due to their complex and symbiotic lifestyles. Against the common belief of haploidy, a recent genomic study found a triploid-like signal in *Letharia*. Here, we used genomic data from a pure culture and from thalli, together with a PCR survey of the MAT locus, to infer the genome organization and reproduction in *Letharia*. We found that the read count variation in the four *Letharia* specimens, including the pure culture derived from a single sexual spore of *L. lupina*, is consistent with haploidy. By contrast, the *L. lupina* read counts from a thallus’ metagenome are triploid-like. Characterization of the mating-type locus revealed a conserved heterothallic configuration across the genus, along with auxiliary genes that we identified. We found that the mating-type distributions are balanced in North America for *L. vulpina* and *L. lupina*, suggesting widespread sexual reproduction, but highly skewed in Europe for *L. vulpina*, consistent with predominant asexuality. Taken together, we propose that *Letharia* fungi are heterothallic and typically haploid, and provide evidence that triploid-like individuals are rare hybrids between *L. lupina* and an unknown *Letharia* lineage, reconciling classic systematic and genetic studies with recent genomic observations.

## Introduction

The way an organism reproduces is one of its most important traits, since patterns of inheritance drastically affect a species ability to adapt and survive over evolutionary time (Bell 1982; Whittle et al. 2011). In Fungi, mating types are key players in reproduction, as they define the sexual identity of an individual at the haploid stage, making it able to recognize and mate with compatible cell types (Kronstad and Staben 1997). Researchers can use the structure of the mating-type (MAT) locus, and the frequency of mating types in natural populations to shed light on the reproductive biology of a species. In particular, the configuration of the MAT locus can inform us about the species’ sexual system.

The most basic MAT configuration occurs in heterothallic ascomycetes, where either of two allelic variants (also referred to as idiomorphs as they represent highly dissimilar sequences that encode different transcription factors) are found at the MAT locus (Butler 2007): one contains a gene encoding a transcriptional activator protein with an α-box domain (referred to herein as *MAT1-1-1*, see Turgeon and Yoder (2000)), and the other contains a gene encoding a high-mobility group (HMG) protein (*MAT1-2-1*). In heterothallic species, successful mating is only possible between haploid individuals of opposite mating types (Glass et al. 1988). By contrast, both idiomorphs are often present in homothallic species, allowing them to mate with itself, and in many cases with any other individual (e.g., Billiard et al. 2012; Gioti et al. 2012). In addition, the idiomorphic regions of many ascomycetes harbour additional MAT genes, which may contribute to the sexual development of the species (e.g., Ferreira et al. 1998; Mandel et al. 2007; Tsui et al. 2013; Dyer et al. 2016; Hutchinson et al. 2016). Moreover, the relative frequency of allelic variants at the MAT-locus in natural populations can be used to investigate the reproductive mode of the species (e.g., Mandel et al. 2007), as an even distribution indicates frequent sexual reproduction while an uneven distribution points towards a relatively high frequency of asexual propagation. Within the class Lecanoromycetes, which includes many lichen symbionts, multiple studies have found a single mating type per sample, indicating a widespread heterothallic sexual system (Singh et al. 2012; Ludwig et al. 2017; Allen et al. 2018; Dal Grande et al. 2018; Pizarro et al. 2019). However, homothallic Lecanoromycetes do exist, such as some members of the genus *Xanthoria s. lat.* (Honegger et al. 2004; Scherrer et al. 2005), emphasizing that taxa must be investigated in a case-by-case manner.

Ascomycete lichen symbionts are usually assumed to grow vegetatively as haploids, as the majority of Ascomycota (Honegger and Scherrer 2008). However, there have been suggestions of alternative genetic compositions present in lichens, such as different nuclear organization, like dikaryosis (n+n), diploidy, and higher ploidy levels (Tripp et al. 2017; Tripp and Lendemer 2018; Pizarro et al. 2019) or as intrathallus variation, in which multiple individuals (genotypes) of the lichen fungal symbiont come together into a single thallus but without cell fusion (Jahns and Ott 1997). The latter phenomenon is also known as “chimeras” or as “mechanical hybrids” (Murtagh et al. 2000; Dyer et al. 2001; Altermann 2004). Commonly, authors make no distinction between different nuclear organizations and the “chimera” scenarios (Honegger et al. 2004; Mansournia et al. 2012). This is partly due to the fact that very little is known about the genetics and composition of thalli, as few species have been successfully grown in isolation from their symbiotic partners (Crittenden et al. 1995; Honegger et al. 2004; Honegger and Zippler 2007). Historically, it has remained extremely difficult to artificially reconstruct the full lichen symbiosis and to mate different individuals in the lab (Bubrick and Galunj 1986; Jahns 1993; Stocker-Wörgötter 2001). However, the advent of next generation sequencing technologies has opened the door to re-evaluate many of our assumptions on lichen biology, including the genetic composition of the dominant ascomycete partner (Tripp et al. 2017; Dal Grande et al. 2018; Armaleo et al. 2019; McKenzie et al. 2020). In particular, Tripp et al. (2017) inferred the ploidy level of seven unrelated fungal lichen symbionts using sequencing data from cultures derived from multiple spores, as well as from thalli containing all symbiotic partners (i.e. metagenomic data). They found evidence of some lineages deviating from haploidy. As compelling as such results may initially seem, such a pattern could have multiple explanations, not all of which involve ploidy changes. Multispore cultures could represent a collection of siblings that would naturally look different from haploid samples. Moreover, it is unclear if lichen fungi have a vegetative incompatibility system in place as other ascomycetes (Dyer et al. 2001; Sanders 2014), which could influence the fusion or lack thereof between germinating spores. Thus, there is no clear expectation as to what would happen in a multispore culture in which different genotypes might meet and undergo cell fusion. Furthermore, the metagenomes studied might include sexual fruiting bodies (apothecia) or nearby tissue, which presumably contains fertilized (dikaryotic or diploid) cells. Hence, general ploidy levels in lichen thalli remain unclear.

In a recent study, McKenzie et al. (2020) used the Oxford Nanopore MinION technology to produce long-read data from whole thalli of two species from the lichen genus *Letharia*: *Letharia lupina* and *Letharia columbiana*. Unexpectedly, they discovered the presence of three *Letharia* genotypes in the metagenomic data of both samples, suggesting that *Letharia* fungal symbionts are triploid-like. They took advantage of the fact that one of the three genotypes within the *L. lupina* sample was markedly different from the other two, allowing them to phase the assembly graph and to produce a highly continuous haploid-like genome assembly. While methodologically convenient, their findings are nothing short of puzzling, since *Letharia* is a genus that belongs to the diverse family Parmeliaceae, whose members are heterothallic and in general assumed to be haploid (Honegger and Zippler 2007; Pizarro et al. 2019). In plant and animal taxa, triploid individuals often have difficulties during meiosis as the chromosomes are unbalanced (Ramsey and Schemske 1998; Tiwary et al. 2005; Köhler et al. 2009), and many higher ploidy populations often persist via asexual reproduction or selfing (Levin, 1975; Mable 2003; Bicknell and Koltunow 2004). The sexual system of *Letharia* has not been studied, but it can be hypothesised that it follows the heterothallic, self-infertile ancestral state. How, then, do triploid or triploid-like fungal symbionts deal with sexual reproduction under heterothallism?

In this study, we explored the reproductive biology of *Letharia.* In their seminal paper, Kroken and Taylor (2001a) studied the diversity within this genus and defined six cryptic, closely related species based on the analyses of 12 loci and their coalescent patterns. That and several subsequent studies have further refined these species hypotheses based on morphological characters, photosynthetic partner lineage, and their assumed main reproductive mode — sexual or asexual (Kroken and Taylor 2001a; Altermann et al. 2014; Altermann et al. 2016). Sexual *Letharia* lineages (referred to as “apotheciate”) produce multiple apothecia that release sexual fungal spores, while asexual lineages (“sorediate”) produce abundant asexual symbiotic propagules also known as soredia. The distinction, however, is not clear-cut as apotheciate species might have isidia, another type of symbiotic asexual propagule, and sorediate species occasionally produce apothecia (Kroken and Taylor 2001a). The global species diversity centre of *Letharia* is found in western North America, where all six lineages of the genus are present. In contrast, only the sorediate species *L. lupina* and *L. vulpina* are known in parts of Europe, Asia and Africa. *Letharia vulpina*, the only representative of the genus in Norway and Sweden, is red-listed in these countries, where it produces apothecia extremely rarely and the genetic variation is low (Högberg et al. 2002; Arnerup et al. 2004; SLU ArtDatabanken 2015; Henriksen and Hilmo 2015).

Here we aimed at collecting *Letharia* samples to represent the diversity of species as previously defined. We gathered genomic data from thalli of four taxa (as metagenomes), as well as from a pure culture derived from a single spore of the species *L. lupina*, and performed PCR surveys of the mating type in additional specimens. With this data in hand, we re-evaluated the ploidy level and verified the sexual system in *Letharia*. We also contrasted patterns of mating type frequencies in different populations of the sorediate species to get insight into the occurrence of sexual vs. asexual reproduction in the species of conservation concern.

## Methods

### Lichen material used in the study

We sequenced the genome and the transcriptome of the pure-cultured *L. lupina* (culture number: 0602M), originating from a single spore from Canada and maintained in the culture collection at Akita Prefectural University (Yamamoto et al. 1998). Likewise, we sequenced both the metagenome and the metatranscriptome of one lichen thallus from each of four *Letharia* taxa: *L. columbiana*, *L. lupina*, *L. ‘rugosa’*, and *L. vulpina*, all collected from Montana, U.S.A. (Table S1, SRA: SRP149293 from Tuovinen et al. (2019)) (fig 1A). In addition, we used 317 *Letharia* thalli for PCR analyses of the mating type from Sweden (N=98; six localities with 1-20 thalli/locality), Switzerland (N=18; one locality), Italy (N=60; six localities with10 thalli/locality) and the U.S.A. (N=138; 14 localities with 1-31 thalli/locality; **Table S1**). All lichen specimens were collected from different trees in order to avoid collecting clones of the individual thalli dispersed on the same tree trunk, except for eight putative *L. gracilis* thalli which we received from B. McCune, all collected from the same tree. In Sweden, only a small part of each thallus (< 3 cm in length) was collected due to the red-listed status of *L. vulpina*. After collection, the specimens were air-dried and stored at –20°C until DNA was extracted. The thalli used for RNA extraction were snap frozen in the field in liquid nitrogen, transferred to the lab and stored at –80°C until further processing. *Letharia* species were identified based on a combination of morphology and internal transcribed spacer (ITS) sequence, as the combination of these characteristics should distinguish the previously described cryptic species (Kroken and Taylor 2001a). In addition, the algal ITS identity was used to assign two sorediate specimens to species following Altermann et al. (2016) (data not shown). We aligned the ITS sequences from all specimens included in this study (GenBank accessions MG645014–MG645052 from Tuovinen et al. (2019)) with the previously published *Letharia* ITS sequences from Kroken and Taylor (2001a, TreeBASE ID # 1126), Altermann et al. (2014, TreeBASE ID # 15485) and Altermann et al. (2016, NCBI PopSet 1072900244) using MUSCLE in AliView v.1.18 (Larsson 2014). Detailed information about the specimens, their collection sites and ITS variants is given in **Table S1**, and the specimens are deposited in the Uppsala University Herbarium (UPS).

**Figure 1.**
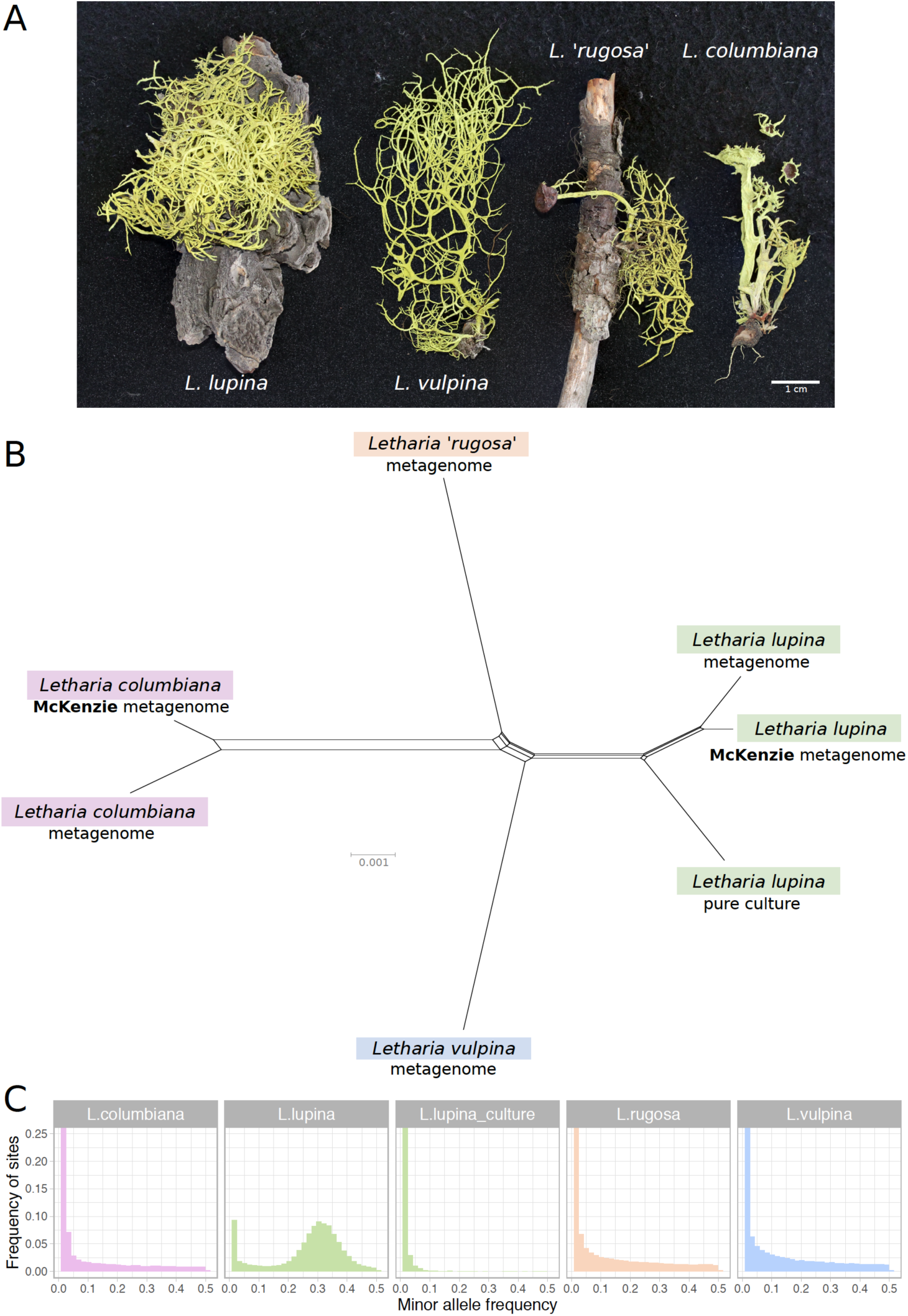
Relatedness and minor allele frequency (MAF) distributions of *Letharia* specimens. **A)** Thalli used for metagenome sequencing in this study. **B)** Unrooted NeighborNet split network based on 680 genes from our samples with genomic or metagenomic data available and those from McKenzie et al. (2020). **C**) Folded MAF distributions of the metagenomes from *Letharia* specimens in (**A**), as well as the whole genome derived from the pure culture of a single *L. lupina* spore. All samples show distributions concordant with haploidy, with the exception of the triploid-like metagenome of *L. lupina*.

### DNA and RNA extraction

For the reference genome and transcriptome sequencing, mycelia of the pure culture of *L. lupina* was placed in a 2 ml safe lock tube with two sterile 3 mm tungsten carbide beads, frozen in a liquid nitrogen and pulverized in a TissueLyzer II (Qiagen) with 25 r/sec for 1 min. In addition, pieces of the vegetative thallus part, carefully avoiding any apothecia or their close proximity, of four *Letharia* used for metagenome and metatranscriptome sequencing and 317 specimens used for PCR screening were physically disrupted as described in Tuovinen et al. 2019. DNA was extracted from both mycelia and lichen tissue by using the DNeasy Plant Mini kit, following the manufacturer’s instructions with a prolonged incubation at 65°C for 1 hr. RNA of the mycelia from the pure culture was extracted with RNeasy Plant Mini Kit (Qiagen) with RLT buffer. RNA from the thalli was extracted with the Ambion Ribopure Yeast Kit (Life Technologies). For detailed description of DNA and RNA extraction of the lichen thalli, see Tuovinen et al. 2019.

### Library preparation and NGS sequencing

Libraries for genome sequencing were prepared from 100 ng of DNA using the TruSeq Nano DNA Sample Preparation Kit (#15041110, rev B). The pure culture was sequenced twice on a split lane of Illumina MiSeq v3 with 300bp paired-end sequencing. The DNA from each thallus was sequenced on 1/5 lane of Illumina HiSeq 2500 with 100bp paired-end sequencing by the SNP&SEQ Technology Platform, Science for Life Laboratory at Uppsala University (Tuovinen et al. 2019).

Libraries for the mRNA sequencing were made by using the Illumina TrueSeq Stranded mRNA Kit (#15031047, rev E) and sequenced with Illumina HiSeq 2500. The transcriptome of the pure culture was sequenced on 1/5 of a lane at the SNP&SEQ Technology Platform, and the metatranscriptomes from the thalli on 1/6 of a lane each at the Huntsman Cancer Institute, University of Utah sequencing core (Tuovinen et al. 2019).

### Quality control and genome and transcriptome assembly

Both the DNA and RNA raw reads were trimmed and quality controlled using Trimmomatic 0.32 (Bolger et al. 2014) (with options ILLUMINACLIP:/adapters.fa:2:20:10:4:true LEADING:10 TRAILING:10 SLIDINGWINDOW:4:15 MINLEN:25 HEADCROP:0). Data quality was inspected using FastQC (Patel and Jain 2012). The draft genome of the pure cultured *L. lupina* was *de novo* assembled from the trimmed reads using SPAdes v. 3.5.0 (Bankevich et al. 2012) with k-mers 21,33,55,77,97,127 and the --careful flag. The raw reads of the metagenomes were assembled with SPAdes using k-mers 55, 75, 85, 95, as preliminary analyses using trimmed reads performed worse (not shown). Likewise, downstream analyses of the *L. lupina* pure culture were done with trimmed reads while the raw reads were used for the metagenomes. A summary of the genome assembly statistics is presented in Table S2 as calculated with QUAST 3.2 (Gurevich et al. 2013). The filtered RNA reads of the pure cultured *L. lupina*, excluding the orphan reads produced after trimming, were assembled using Trinity 2.0.6 (Grabherr et al. 2011; Haas et al. 2013).

### Genome characterization of *L. lupina*

As the long-read *L. lupina* assembly from McKenzie et al. (2020), hereafter referred to as the McKenzie *L. lupina* assembly, was derived from an entire lichen thallus, we compared it to the SPAdes assembly of the pure-cultured *L. lupina*. We produced whole genome alignments with the NUCmer program from the MUMmer package v. 4.0.0beta2 (Kurtz et al. 2004) with options -b 2000 -c 200 --maxmatch. This analysis revealed only small alignments (less than 11 kbp) between the contig 1 of the McKenzie *L. lupina* assembly and multiple scaffolds from the SPAdes assembly of the pure-cultured *L. lupina*. To evaluate if the contig 1 of the McKenzie *L. lupina* assembly really is part of the genome of the ascomycete and not other symbiotic partners, we explored the repetitive content of both assemblies with RepeatMasker v. 4.0.8 (http://www.repeatmasker.org/) and the repeat library from McKenzie et al. (2020) with parameters -a -xsmall -gccalc -gff -excln.

### Phylogenetic analyses of marker genes

Alignments were produced with MAFFT version 7.245 (Katoh and Standley 2013) and options - -maxiterate 1000 --retree 1 --localpair. We used the program RAxML v. 8.0.20 (Stamatakis 2014) to infer a Maximum Likelihood (ML) tree of the previously published *Letharia* ITS variants along with variants discovered in our data. Likewise, we produced ML trees from the idiomorph sequences (MAT1-1 or MAT1-2) as above but partitioning by gene and intergenic regions. Branch support was assessed with 1000 bootstrap pseudo-replicates. The specimens with unique ITS variants were putatively assigned to species based on the ML tree, previously published informative SNPs that can separate between *L. vulpina* and *L. lupina*, and previously described morphological characters (Kroken and Taylor 2001a; McCune and Altermann 2009; Altermann et al. 2014; Altermann et al. 2016).

In order to place our genomic samples within the diversity of the genus, we extracted the sequence from 11 markers used in previous *Letharia* studies (ribosomal ITS, the 18S intron, chitin synthase I, and the anonymous loci 11, 2, 13, DO, CT, BA, 4, and CS) from the haploid individuals (see below) and fused them with the concatenated alignment from Altermann et al. 2016 (TreeBASE study ID S18729). We then inferred a ML topology with IQ-TREE v. 1.6.8 (Nguyen et al. 2015). Branch support was estimated with 100 standard bootstraps. Rooting was arbitrarily done with the well-supported *L. columbiana* (*L. ‘lucida’*) lineage.

### Network analysis of genomic data

Previous population studies using few markers have shown that the *Letharia* species are very closely related (Kroken and Taylor 2001a; Altermann et al. 2014; Altermann et al. 2016). In order to approximate the genetic distance and relationships between the *Letharia* individuals for which we had whole-genome data available, we used as reference the genome annotation of the McKenzie *L. lupina*. We randomly selected 1000 gene features, extracted their corresponding sequences from the McKenzie *L. lupina* assembly (including introns), and used them as queries in BLAST searches performed on the SPAdes assembly of our four metagenomes and the *L. lupina* pure culture. We also included the long-read assembly of the *L. columbiana* individual from McKenzie et al. (2020), hereafter referred to as the McKenzie *L. columbiana*. We filtered the BLAST output to include only hits that had at least a 250bp-long hit alignment and 90% identity to the query. We extracted the sequence corresponding to the surviving hits and aligned them using MAFFT version 7.458 (Katoh and Standley 2013) with options --adjustdirection --maxiterate 1000 --retree 1 --localpair. With the aim of approximating single-copy orthologs, only those alignments that contained a single hit for all samples were retained. The resulting 680 genes were concatenated into a matrix of 1,184,497 bp from which 35,458 bp were variable. We used the matrix as input for SplitsTree v. 4.14.16, build 26 Sep 2017 (Huson and Bryant 2006) to produce an unrooted split network with a NeighborNet (Bryant and Moulton 2004) distance transformation (uncorrected distances) with an EqualAngle splits transformation and excluding gap sites.

### Inferring ploidy levels based on the minor allele frequency (MAF) distribution

As the McKenzie samples were inferred to be triploid-like (McKenzie et al. 2020), we determined the ploidy levels of our metagenomes and the pure culture by calculating the distribution of the minor (i.e. less common) allele in read counts of biallelic SNPs (Yoshida et al. 2013; Ament-Velásquez et al. 2016). Normally, a diploid individual should have a minor allele frequency (MAF) distribution around 0.5, while in a triploid individual the mode should be around 1/3. A tetraploid individual is expected to have two overlapping curves, one centered around 0.25 and another around 0.5. The width of the curves is proportional to factors like coverage variance, mapping biases, and hidden paralogy (Heinrich et al. 2011). In addition, any given sample is expected to have a large number of very low frequency alleles due to sequencing errors. As a result, a haploid sample should only exhibit a curve corresponding to the sequencing errors. Note that a caveat of this approach is that the haploid signal cannot be distinguished from an extremely homozygous higher-ploidy level individual, as these will show few heterozygous sites. Moreover, a diploid or higher ploidy signal in MAF distributions could correspond to a heterokaryon condition (n+n, n+n+n, etc), which strictly speaking is composed of haploid nuclei. Hence, “ploidy” is used here in a broad sense to include genetically different heterokaryons.

To produce MAF distributions, we mapped the Illumina reads of all of our metagenomes to the pure-cultured *L. lupina*, which should exclude most reads from other symbionts, using BWA v. 0.7.17 (Li and Durbin 2010) with PCR duplicates marked by Picard v. 2.23.0 (http://broadinstitute.github.io/picard/). To call variants we used VarScan2 v. 2.4.4 (Koboldt et al. 2012) with parameters --p-value 0.1 --min-var-freq 0.005 to ensure the recovery of low frequency alleles and no further influence of the variant caller on the shape of the MAF distribution. Only biallelic SNPs from contigs larger than 100 kbp and with no missing data were used. We further filtered out all SNPs overlapping with repeated elements annotated above with RepeatMasker, and modified the output of VarScan2 to be compliant with the vcf format (https://samtools.github.io/hts-specs/). Based on the general coverage levels of all samples, we excluded variants with coverage above 300x or below 50x. These high quality SNPs were processed with the R packages vcfR v. 1.10.0 (Kamvar et al. 2015; Knaus and Grünwald 2017), dplyr v. 0.8.0.1 (Wickham et al. 2018), ggplot2 v. 3.1.1 (Wickham 2016), and cowplot v. 1.0.0 (Wilke 2019). MAF distributions were scaled such that the area of all bars amount to 1. A full Snakemake v. 5.19.2 (Köster and Rahmann 2012) pipeline is available at https://github.com/johannessonlab/Letharia/tree/master/LichenPloidy.

### Measuring distance of the subgenomes within the *L. lupine* metagenome

Given the presence of two distinguishable genotypes in the metagenome of *L. lupina* (see Results), we calculated a relative measure of distance of each subgenome with respect to the other *Letharia* lineages (*L. columbiana*, *L. rugosa*, and *L. vulpina*). We assumed that the pure-cultured *L. lupina* corresponds to the “true” *L. lupina* genotype, and assigned all sites in the *L. lupina* metagenome with identical genotype to the pure culture as “lupina”, while all alternative alleles were treated as “other”. For each *Letharia* lineage, we counted the proportion of sites that were not identical to either the “lupina” or the “other” variant. We made a further distinction between areas along the scaffolds that were polymorphic (having both “lupina” and “other” alleles), or fixed for either allele within the *L. lupina* metagenome. As input we used the filtered SNPs produced above but we discarded all sites with MAF < 0.2 (*L. lupina* metagenome) or MAF < 0.9 (haploid samples) under the assumption that they correspond to sequencing errors. Only sites with data from all samples were considered.

### Identifying scaffolds of the genomes containing the MAT locus

We performed searches with the BLAST 2.2.31+ suit (Camacho et al. 2009) of the MAT-gene domains (HMG and α; see Martin et al. 2010) in the assembly of the *L. lupina* pure culture. As queries we used the corresponding gene sequences of *Polycauliona polycarpa* (*Xanthoria polycarpa*; Lecanoromycetes, Genbank accession numbers AJ884598 and AJ884599). Furthermore, we searched for *APN2* (encoding a basic endonuclease/DNA lyase) and *SLA2* (encoding a protein that binds to cortical patch actin) since these genes are typically flanking the MAT genes in Pezizomycotina (Heitman et al. 2007). As queries we used the ortholog clusters (OG5 126768 and OG5 129034 for APN2 and SLA2, respectively) retrieved from OrthoMCL-DB (Chen et al. 2006) and restricted to Pezizomycotina. Finally, we used the MAT locus retrieved from the *L. lupina* pure culture assembly as a query to perform BLAST searches against the assemblies of the metagenomes (*L. lupina*, *L. vulpina*, *L. columbiana*, and *L. ‘rugosa’*).

### Annotating the MAT locus

We used three *ab initio* gene predictors to annotate *Letharia* scaffolds containing the MAT locus: SNAP release 2013-02-16 (Korf 2004), Augustus v. 3.2.2 (Stanke and Waack 2003) and GeneMark-ES v. 4.32 (Lomsadze et al. 2005; Ter-Hovhannisyan et al. 2008). All gene predictors were trained for the *Letharia* genomes as described in the Appendix.

We used STAR v. 2.5.1b (Dobin et al. 2013) to map filtered RNAseq reads to the entire genome assembly of the pure-cultured *L. lupina*, or to the relevant scaffold extracted from the metagenomes. Cufflinks v. 2.2.1 (Trapnell et al. 2010) was used to create transcript models from which ORFs were inferred using TransDecoder v. 3.0.1 (Haas et al. 2013). The ortholog sequences of *APN2* and *SLA2* of *Aspergillus oryzae* (OrthoMCL-DB: RIB40_aory) and the MAT genes of *Polycauliona polycarpa* were aligned to the *Letharia* scaffolds using Exonerate v. 2.2.0 (Slater and Birney 2005) with the options --exhaustive TRUE --maxintron 1000. Using all these sources of evidence, we created weighted consensus gene structures using EVidenceModeler (EVM) v. 1.1.1 (Haas et al. 2008) that were subject to manual curation with Artemis release 16.0.0 (Rutherford et al. 2000).

To determine the identity of auxiliary MAT genes found in *Letharia*, we used the protein sequences of *L. rugosa* (MAT1-1) and *L. columbiana* (MAT1-2) to conduct BLASTp and DELTA-BLAST searches in the NCBI Genbank database (Boratyn et al. 2012) (searches conducted the 29th of April of 2020), as well as HMMER v. 3.3 (www.hmmer.org; Eddy 2011) with the uniprotrefprot database, version 2019_09. We also compared the sequences with the protein models provided by Pizarro et al. (2019) for multiple lecanoromycete species. In addition, we re-annotated the scaffold with the MAT locus of the species *Leptogium austroamericanum* and *Dibaeis baeomyces* (Pizarro et al. 2019), as well as the MAT1-2 idiomorph of *Cladonia grayi* (accession number MH795990; Armaleo et al. 2019) using the pipeline above in order to confirm their gene order and composition. We found that the protein model of the MAT1-1 auxiliary gene as provided by Pizarro et al. (2019) is relatively variable at the C’ terminus, which could be explained by inconsistencies in the gene annotation process in the absence of sources of evidence additional to the *ab initio* prediction (e.g. RNAseq data), or by true biological differences. Hence, only the first exon was analysed further with IQ-TREE. Likewise, we only used the first and second exon of the MAT1-2 auxiliary gene for an IQ-TREE analysis.

### PCR screening of the mating type and additional annotation of the idiomorphs

We designed idiomorph specific primer pairs to screen for the mating-type ratios in different populations based on the annotation of the MAT-loci from the genomes (MAT1-1: MAT11L6F and MAT11L6kR, MAT1-2: MAT12LF1 and MAT12LR1; Table S3). Each sample was amplified with primer pairs for both idiomorphs and the presence/absence of the PCR-product of each idiomorph was determined by using agarose gel electrophoresis. A subsample of each idiomorph was sequenced to ensure the amplification of the correct target sequence with the PCR primers. We tested whether the distribution of mating types in the sorediate species (*L. lupina* and *L. vulpina*) in U.S.A., the Alps and Sweden follows the 1:1 ratio expected for a population that regularly reproduces sexually by using an exact binomial test with 95% confidence interval.

In order to sequence and annotate both idiomorphs for each *Letharia* species, we selected specimens of the same species (based on ITS sequence information) as the individual from which the genomic data originated, but of the opposite mating type, and amplified a *circa* 3800 bp long part of the MAT locus using primers LMATF2 and LMATR1 and APN2LF and SLA2LR (Table 1). The same procedure was done for *L. ‘barbata’* and the putative *L. gracilis* specimens. The entire MAT locus was then sequenced with primers designed for *Letharia* based on the genome annotations (Table 1). The PCR-obtained sequences of the idiomorphs were annotated using the trained *ab initio* gene predictors and the alignments (with Exonerate) of the final gene models. We define the idiomorphic part of the MAT locus as the whole region between *APN2* and *SLA2* that lack sequence similarity between individuals of different mating types.

## Results

### *Letharia* species in the study

Based on morphology and ITS sequence variation, we assigned the collected 322 specimens to five of the six previously recognized *Letharia* species (Kroken and Taylor 2001a; Altermann 2009; Altermann et al. 2016), including those used for metagenome sequencing (see Appendix). While traditionally-used markers generally provide little resolution to the *Letharia* species complex (fig. S1 and S2; Altermann et al. 2014; Altermann et al. 2016), a NeighborNet split network of 680 genes extracted from all (meta)genome assemblies confirmed clustering by species for these samples (fig. 1B).

### High correspondence between our pure culture assembly and the McKenzie assembly of *L. lupina*

The McKenzie *L. lupina* assembly was produced by manual curation of metagenomic long-read data and is one of the best genome assemblies for an ascomycete lichen symbiont currently available (McKenzie et al. 2020). However, it could still contain sequences from other symbiotic partners. As a sanity check, we compared the *L. lupina* pure culture assembly generated by us to the McKenzie *L. lupina* assembly and found that all contigs, except contig 1, have good correspondence between the two. There are very few obvious translocations or inversions, which might be either assembly artifacts or biological differences between samples (fig. S3). Conversely, contig 1 only produced very small (< 11 kbp) alignments to multiple scaffolds of the *L. lupina* pure culture assembly (fig. S4). This contig is the largest in the McKenzie *L. lupina* assembly (> 3 Mbp), with a size comparable to the smallest chromosomes of well-studied ascomycetes like *Podospora anserina* (Espagne et al. 2008). Remarkably, contig 1 is composed of 87% repetitive elements, in stark contrast with most other contigs that harbor around 30% repetitive elements (fig. S4). We reasoned that if the repetitive elements present in contig 1 also occur in the other contigs of either the McKenzie *L. lupina* or the pure-cultured *L. lupina*, then this large contig is indeed part of the *L. lupina* genome and not of another symbiont. Accordingly, we found that the five most abundant repeats of contig 1 are present elsewhere in both assemblies (fig. S6 and fig. S7), and were probably not recovered as such in the pure culture assembly due to the limitations of short-read technology. Hence, we conclude that the McKenzie *L. lupina* assembly does reflect the genome of the ascomycete *L. lupina* and the observation of a triploid-like signal is not an artifact of additional unrelated fungal sequences in their assembly.

### Genomic and metagenomic datasets show haploid-like patterns except for the metagenome of *L. lupina*

As the results of McKenzie et al. (2020) showed a triploid-like signal for *L. lupina* and *L. columbiana*, we inferred the ploidy levels in our genomic and metagenomic samples by producing MAF distributions using the *L. lupina* pure culture as reference for variant calling. As expected under haploidy, all but one sample show an exponential decay of low frequency variants produced by sequencing errors, including the *L. lupina* pure culture (fig. 1C). The exception to the haploid pattern is our *L. lupina* metagenome, which has a clear peak centered around ⅓, consistent with triploidy. Note that the exponential decay of the variants of the *L. lupina* pure culture is steeper than for the other samples. This difference results from higher coverage and longer reads, as the *L. lupina* pure culture was sequenced with MiSeq, as opposed to HiSeq for other samples, leading to better read mapping. In addition, its own genome was used as reference for mapping, minimizing the variance further. We obtained similar results using the McKenzie *L. lupina* as reference (data not shown).

### *Letharia* has a heterothallic MAT-locus architecture

Having established that most metagenomes and the pure-cultured *L. lupina* are most likely haploid, we proceeded to characterize the MAT locus to give us an indication of the sexual system in *Letharia*. The MAT locus with the flanking genes *APN2* and *SLA2* was found within a single scaffold for the pure-cultured *L. lupina* (submitted to GenBank under the accession number MK521630) and from the metagenomes of *L. columbiana*, *L. ‘rugosa’,* and *L. vulpina* (GenBank numbers MK521629, MK521632 and MK521631, respectively). Only one mating-type idiomorph was recovered in each assembled genome, suggesting that *Letharia* is heterothallic.

The genome of the pure-cultured *L. lupina* and the metagenomes of *L. vulpina*, and *L. ‘rugosa’* contained the MAT1-1 idiomorph (with the α domain of the *MAT1-1-1* gene), while the MAT1-2 idiomorph (with the HMG domain of the *MAT1-2-1* gene) was found in the metagenome of *L. columbiana*. Fig. 2 shows the gene content of the MAT1-1 idiomorph as represented by *L. ‘rugosa’*, and the MAT1-2 as represented by *L. columbiana*. Notably, alignments with Exonerate revealed the remnants of the last exon of the *MAT1-1-1* gene in the idiomorph MAT1-2 (the * in fig. 2B). Albeit homologous, this piece of exon is highly diverged from the *MAT1-1-1* in *Letharia* MAT1-1, as revealed by the percentage of pairwise identity along the idiomorphs (fig. 2C). This same pattern was observed in the species of the genus *Cladonia* (family Cladoniaceae) by Armaleo et al. (2019), suggesting that the pseudogenized exon might be ancestral to both families. In addition, a smaller ORF was predicted weakly based on transcriptomic data of *L. columbiana* downstream of *APN2* in both idiomorphs (as *lorf*, from *Letharia* open reading frame, in fig. 2; see also fig. S8-S11). Likewise, the transcriptomic data revealed an antisense transcript for the *MAT1-1-1* gene in all species (fig. 2 and fig. S8-S11). See Appendix for more details.

**Figure 2.**
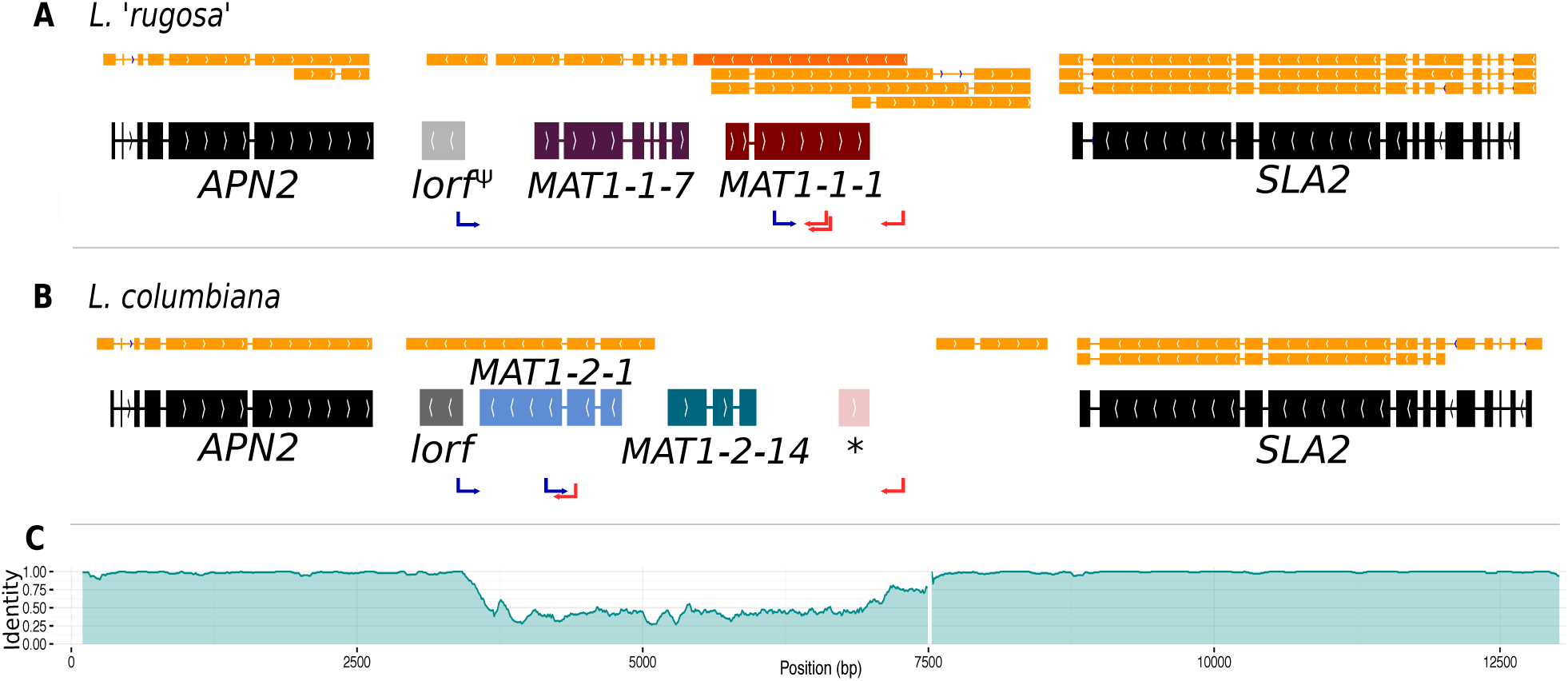
The structure of the MAT idiomorphs in the genus *Letharia*. The typical gene content of the MAT1-1 idiomorph as exemplified by *L. ‘rugosa’* (**A**) is contrasted to the MAT1-2 idiomorph of *L. columbiana* (**B**). The final gene models with intron-exon boundaries are shown along with the transcript models (orange). In (**A**) the antisense transcript of the *MAT1-1-1* gene is marked with a darker orange colour. Notice that the *lorf* gene is pseudogenized in *L. ‘rugosa’* (*lorf^Ψ^*). In (**B**) a star (*) shows the location of a sequence homologous, but highly divergent, to the last exon of the *MAT1-1-1* gene. The pairwise identity between idiomorphs of the MAT locus is plotted in (**C**) using overlapping windows of 100 bp (incremental steps of 10 bp along the sequence), showing the drop in identity. The white gap towards the 3’ end of the area of low identity corresponds to a track of 129 bp in the MAT1-2 idiomorph of all species. The positions of forward (blue) and reverse (red) primers used for PCR screening are marked with arrows.

We further obtained the sequence of the MAT region from additional 46 thalli by PCR, which corresponded to either the MAT1-1 or MAT1-2 idiomorph for all species (except *L. cf. gracilis*, for which all samples had a MAT1-2 idiomorph). As the idiomorphs are very similar between species (fig. S12), we can use the sequence comparison between *L. rugosa* and *L. columbiana* to estimate the size of the non-recombining region in *Letharia* by examining the level of divergence between genomic sequences of MAT1-1 and MAT1-2 (fig. 2C). Towards *APN2*, the identity between sequences drops from 98%-99% (overlapping windows of 100 bp with incremental steps of 10 bp) to on average 48%, marking the beginning of the idiomorphic region. On the 3’ end, towards *SLA2*, a plateau region of around 70% identity covers approximately 500bp. The sequence identity reaches this plateau of intermediate values just before the end of the idiomorph, and it is followed by an indel of 129 bp present in MAT1-2, but not in the MAT1-1, for all *Letharia* species. We consider the 3’-end of this indel the boundary for the 3’-end of the idiomorph. Using these boundaries, we estimate that the size of each idiomorph of *Letharia* is around 3.8 kbp.

### The auxiliary genes present in *Letharia* MAT idiomorphs are ancestral to the Lecanoromycetes

We found that both idiomorphs of the MAT locus of each *Letharia* species harbour additional ORFs (one in each idiomorph) designated herein as *MAT1-1-7* and *MAT1-2-14* (fig. 2). The nomenclature of these additional genes is contentious and discussed extensively in the Appendix.

Pizarro et al. (2019) reported a number of ORFs associated with the lecanoromycete idiomorphs, but it was not clarified whether these so-called “auxiliary MAT genes” are homologous across lichen symbionts or other fungal taxa, making comparisons difficult. Hence, we used protein sequence and synteny comparisons to define homology of these genes. We found that most, but not all, Lecanoromycetes have the same auxiliary gene in the MAT1-1 idiomorph as *Letharia* (i.e., *MAT1-1-7*; left panel of fig. 3) and at the same location (data not shown). In addition, this ORF seems to be homologous to a gene predicted for members of the class Eurotiomycetes (left panel of fig. 3). By contrast, the auxiliary genes present in the MAT1-1 idiomorph of *Leptogium austroamericanum* (Collemataceae, Lecanoromycetes) and *Dibaeis baeomyces* (Icmadophilaceae, Lecanoromycetes) (Pizarro et al. 2019) are not related to *MAT1-1-7*, and are located in a different position (between *MAT1-1-1* and *SLA2*).

**Figure 3.**
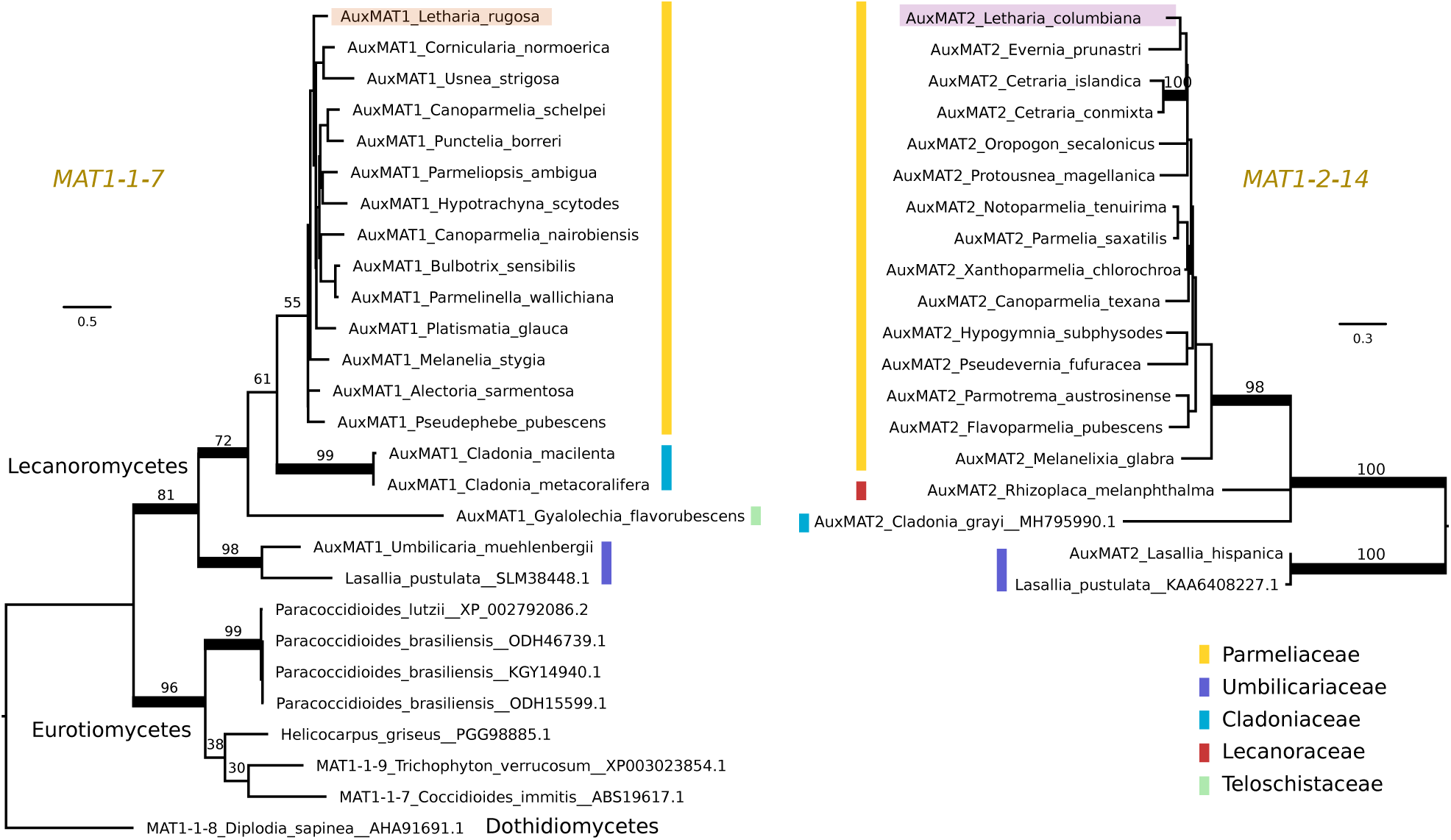
Maximum likelihood trees of the auxiliary genes *MAT1-1-7* (left) and *MAT1-2-14* (right). Colored vertical bars mark the different families within the lichen-forming class Lecanoromycetes, and the *Letharia* sequences are highlighted. *MAT1-1-7* is found outside Lecanoromycetes, while *MAT1-2-14* seems unique for this class. Notice that the homologous genes of *MAT1-1-7* in *Trichophyton verrucosum* and *Diplodia sapinea* have been named differently by other authors. Branches with bootstrap values above 70% are thickened. Branch lengths are proportional to the scale bar (amino acid substitution per site).

For the *Letharia* auxiliary gene in the *MAT1-2* idiomorph (*MAT1-2-14* in fig. 2), we also found homologs in other Lecanoromycetes (right panel of fig. 3), but no homolog outside of Lecanoromycetes. This gene was not identified by Armaleo et al. (2019) in *Cladonia*, but our prediction pipeline recovered it in the *Cladonia grayi* genome (right panel of fig. 3) in the expected position.

Taken together, our data shows that the general idiomorph architecture of *Letharia* is conserved at least between Parmeliaceae, Cladoniaceae, and Umbilicariaceae (fig. 3). Given the presence of *MAT1-1-7* in a member of Teloschistaceae, and of *MAT1-2-14* in a member from Lecanoraceae (fig. 3), potentially the general structure of the idiomorph is the same also in these families.

### The allele frequencies along the genome of the *L. lupina* metagenome suggest hybridization

To explore the triploid-like signal in our *L. lupina* metagenome, we mapped its reads to the pure culture assembly. We assigned the SNPs used for the MAF distribution as either matching the pure culture (“lupina” allele) or having an alternative allele (“other”) and produced an unfolded allele frequency distribution of the metagenome sample (fig. 4). We found that the allele frequency distribution of the “lupina” allele is not identical to that of the “other” allele. In fact, there seems to be a tendency for the “lupina” allele to be in higher frequency (⅔ or 1). Hence, we plotted the distribution of the “other” allele along the pure culture assembly and found that, while there are large regions polymorphic at ⅓ and ⅔ frequencies, there are also areas of loss of heterozygosity (LOH), mostly in favor of the “lupina” allele (fig. 5). The observed genome-wide MAF distribution patterns suggests that there are three haplotypes within the *L. lupina* metagenome. Two of these haplotypes are more similar to each other, as reported for the McKenzie *L. lupina* long-read data, and they correspond to the pure culture *L. lupina*. Such signal is highly suggestive of hybridization between *L. lupina* and another *Letharia* lineage, presumably leading to triploidy (i.e. allopolyploidization) or to a triploid-like condition (n+n+n or 2n+n).

**Figure 4.**
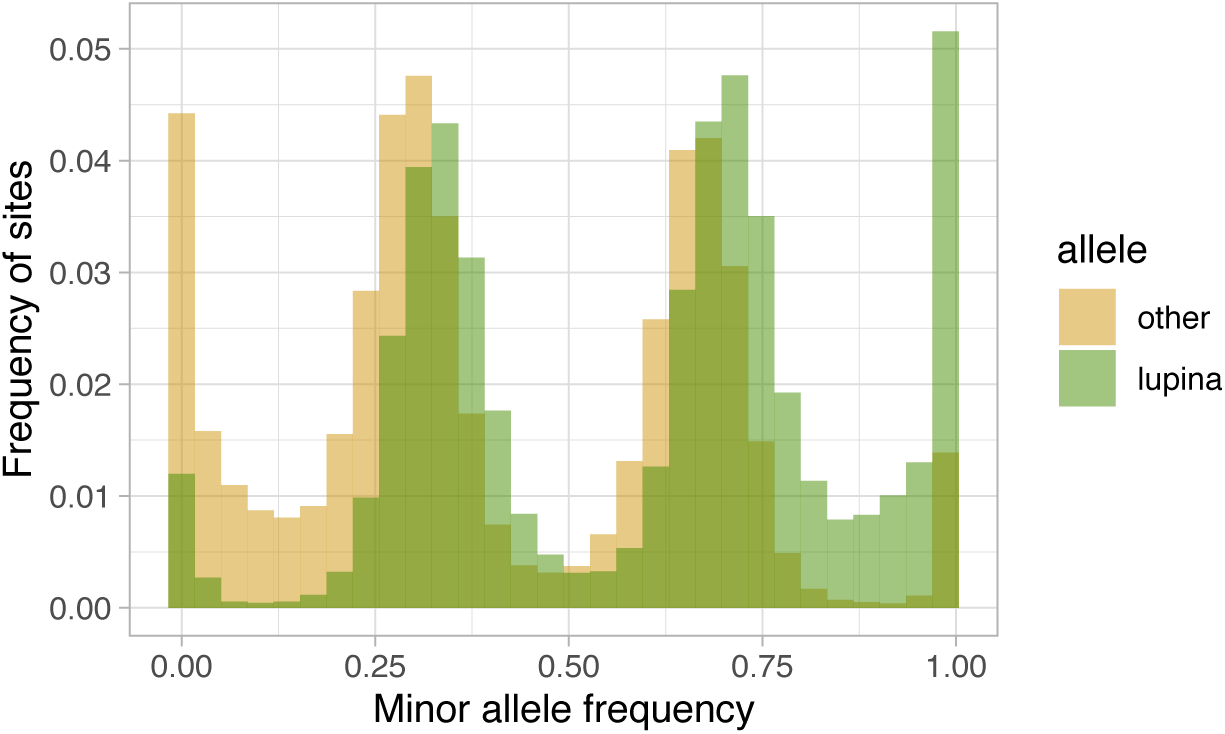
Unfolded allele frequency distribution of the *L. lupina* metagenome, with alleles classified as either identical to that of the *L. lupina* pure culture (“lupina”) or an alternative allele (“other”).

**Figure 5.**
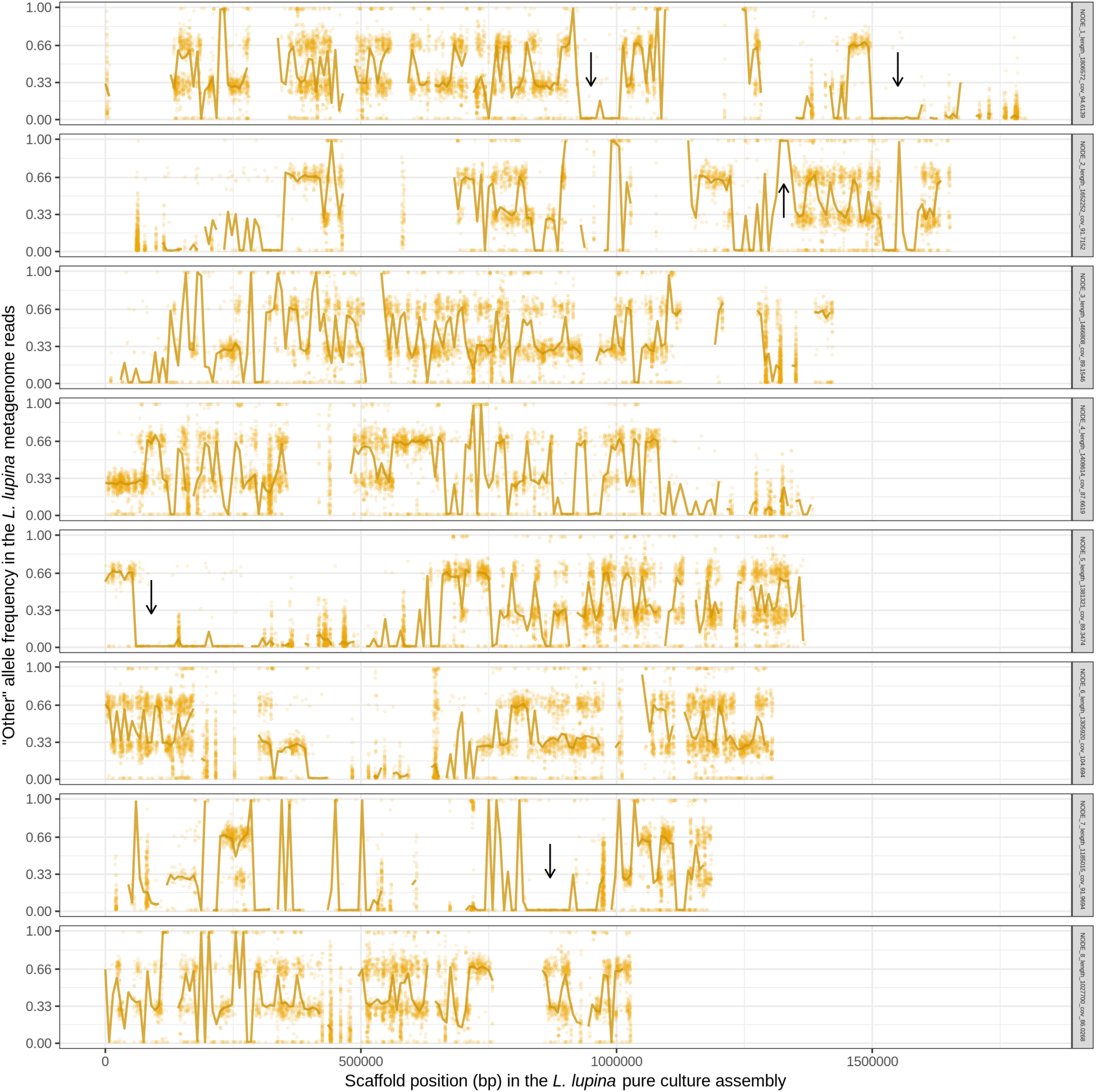
Frequency of the “other” (alternative) allele in the reads of the *L. lupina* metagenome when mapped to the *L. lupina* pure culture assembly (only the eight largest scaffolds are shown). Golden points represent the raw allele frequency at each site, while the solid line connects the median allele frequency of non-overlapping 7.5 kb-long windows. Sites and windows overlapping with repetitive elements were discarded (missing data). Representative areas with long tracks of loss of heterozygosity (LOH) are marked with arrows.

The reads of the *L. lupina* metagenome that map to the scaffold containing the MAT locus also maintain a ⅔ proportion in favor of the “lupina” allele (fig. 6). BLAST searches revealed that our *L. lupina* metagenome contains both MAT idiomorphs. As expected, the coverage of the MAT1-1 idiomorph is around ⅔ of the mean (fig. S13), confirming that two haplotypes are MAT1-1 and the other haplotype is MAT1-2. In addition, we found both idiomorphs in the raw reads of the McKenzie *L. columbiana* metagenome, but not in the McKenzie *L. lupina* one, in which we only found MAT1-1. Thus, the triploid-like *Letharia* thalli can have either both or a single mating type.

**Figure 6.**
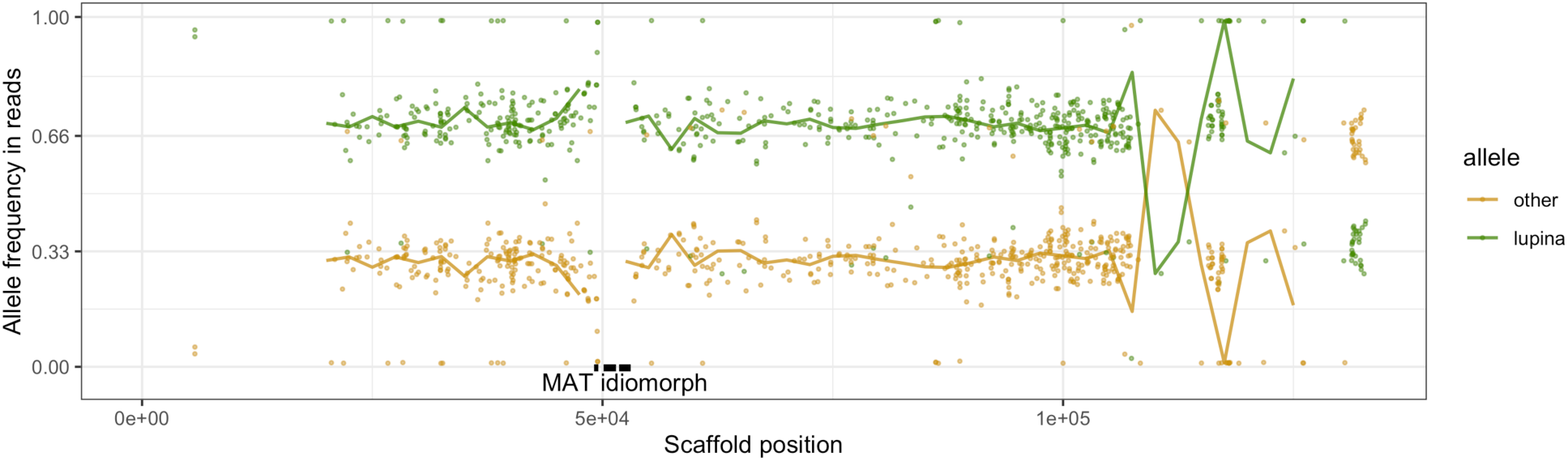
Frequency of the “lupina” and the “other” (alternative) allele in the reads of the *L. lupina* metagenome along the scaffold containing the MAT locus in the *L. lupina* pure culture assembly. Points represent the raw allele frequency at each site, while the solid lines connect the median allele frequency of non-overlapping 2.5 kb-long windows. Sites and windows overlapping with repetitive elements were discarded (missing data). The genes within the MAT1-1 locus are marked with black rectangles.

In an effort to determine the unknown parental lineage, i.e., the origin of the “other” allele, we compared the “lupina” allele and the “other” allele with their homologs in *L. columbiana*, *L. ‘rugosa’* and *L. vulpina*. Surprisingly, in areas where the metagenome remains polymorphic, the “other” allele is twice as divergent from any other known *Letharia* lineage as the “lupina” allele (left panel of fig. 7). Specifically, ∼31% of the “lupina” alleles are different from any other sampled *Letharia*, while the “other” allele is different ∼69% of the time. This finding suggests that the unknown parental lineage is considerably more divergent than any of the sampled species. Surprisingly, this pattern is inverted at sites where LOH occurred in favor of the “other” allele (middle of fig. 7), which suggests that the *L. lupina* pure culture itself (used as reference) has some tracks of highly divergent regions introgressed from the unknown *Letharia* lineage. Notably, at positions where LOH occurs in favor of the “lupina” allele, the proportion of different sites is intermediate in value (right of fig. 7). This might be the result of averaging sites of different ancestry (either “lupina” or “other”) that cannot be recognized as such because they are also introgressed in the reference. Moreover, *L. columbiana* has a higher proportion of sites that are different from the “lupina” allele compared to *L. ‘rugosa’* and specially *L. vulpina*. This pattern might reflect the fact *L. columbiana* has a slightly higher genetic distance to *L. lupina* than the other species sampled here (fig. 1; fig. S2), but it raises the question as to why such a difference is not observed in the other two categories of sites.

**Figure 7.**
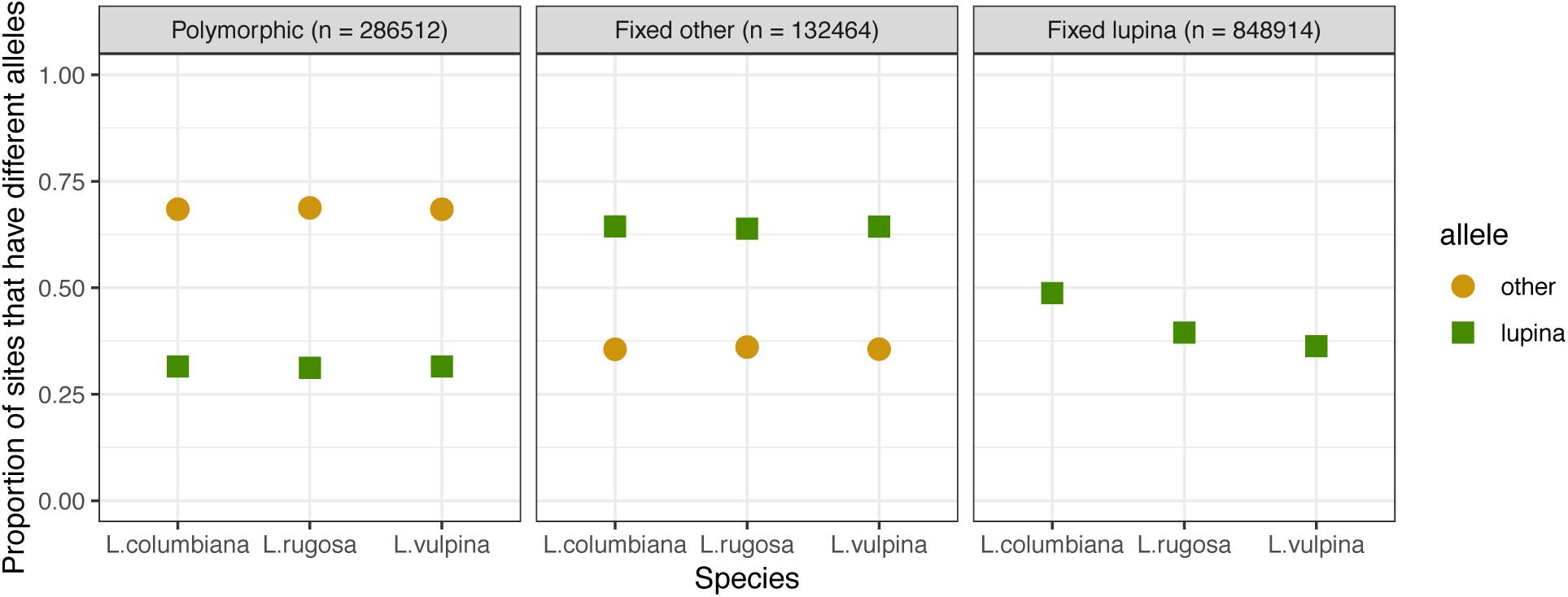
Relative genetic distance between the alleles present in the *L. lupina* metagenome (“lupina” or “other”) and the sampled *Letharia* species: *L. columbiana, L.* ‘*rugosa*’ and *L. vulpina*. All the genotyped sites can be classified as those that are polymorphic in the *L. lupina* metagenome (that is, both the “lupina” or “other” alleles are present), or as sites that are fixed for either the “other” or the “lupina” allele (sites with loss of heterozygosity). If the subgenomes within the *L. lupina* metagenome belong to the same species, the proportion of sites that have different alleles from those present in the sampled *Letharia* species should be roughly the same for both the “other” and “lupina” alleles. However, such a pattern is not observed, suggesting that the “other” allele originates from an unknown, highly divergent *Letharia* lineage. n: number of genotyped sites.

### Regional distribution of mating types in sorediate species shows variation in reproductive mode

We amplified a single idiomorph in all specimens collected with two exceptions: in one *L. vulpina* from Italy and one *L. lupina* from Switzerland we amplified both idiomorphs (**Table S1**). With the aim to investigate the primary reproductive mode of the sorediate species (*L. lupina* and *L. vulpina*), we characterized the regional occurrence of their mating types. In the U.S.A., even when only a few individuals of each species were collected from the same locality, both mating types were sampled. The mating-type ratio in the U.S.A. (when all American samples of a species were treated as one region) of neither sorediate species deviated from a 1:1 ratio (exact binomial test with 95% confidence interval: N=85; *p*=0.8 for *L. lupina* and N=13; *p*=1 for *L. vulpina*). Furthermore, both mating types were present in *L. lupina* in the Alps and the mating-type ratio in these populations did not deviate from 1:1 (N=18; *p*=0.2). However, the ratio of mating types in *L. vulpina* is significantly skewed from a 1:1 ratio (N=59; *p=*1.8e-10) in the Alps, and in Sweden only one mating-type was found in 98 samples (fig. 8, **Table S1**). These findings suggest that sorediate species reproduce sexually in U.S.A, but that *L. vulpina* reproduces sexually at very low frequency in Europe.

**Figure 8.**
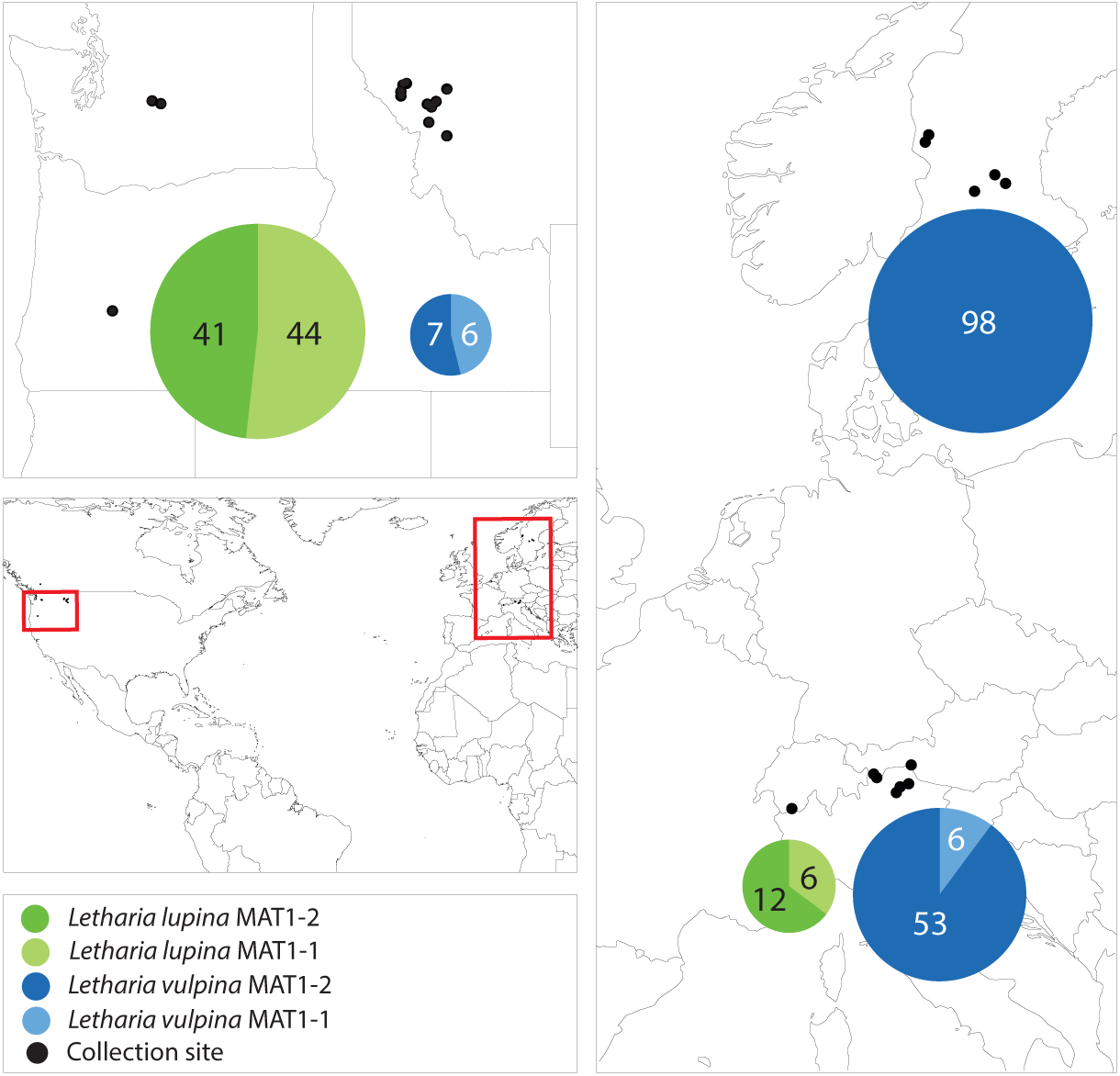
A map showing the collection sites and mating-type distribution in the sorediate species in the U.S.A., the Swiss and Italian Alps and Sweden. The numbers correspond to the number of sampled thalli (n), and the areas of the pie charts are proportional to n.

## Discussion

Our combined approach of next-generation sequencing from representative samples, along with regional PCR surveys of the mating types, has allowed us to revise key aspects of the reproductive biology of *Letharia*. Specifically, we were able to determine haploidy in most genomic samples and a triploid-like signal in one, we revealed conserved heterothallism across the genus, and we uncovered differences in mating type frequencies between North American and European specimens of sorediate species. McKenzie et al. (2020) qualified the genome architecture of *Letharia* as “enigmatic”, referring to the triploid-like nature of their two samples. Here we further point out the unexpectedly high content of repeated elements in the contig 1 of their *L. lupina* assembly. Future research should aim at clarifying if this contig is a core feature of the *Letharia* genome, or if it represents some form of accessory or B chromosome, which typically have a high repeat content (Bertazzoni et al. 2018; Ahmad and Martins 2019).

In contrast to previous studies addressing ploidy levels (Tripp et al. 2017; McKenzie et al. 2020), we made use of a pure culture derived from a single spore, which removes confounding factors related to the presence of other unrelated fungi within the thallus, and to unknown effects of growing alongside siblings. In effect, our pure culture represents the immediate state after sexual reproduction in *L. lupina*. The fact that the metagenomes of *L. columbiana*, *L. ‘rugosa’*, and *L. vulpina* are also haploid further suggests that a single spore is enough, with symbionts, for the development of a lichen thallus, and that a *Letharia* lichen is not formed from a collection of spores as it has been suggested for other species (Jahns and Ott 1997; Murtagh et al. 2000). Crucially, in their classic study, Kroken & Taylor (2001b) separately genotyped ascomata and maternal thalli from four specimens of *L. gracilis* and two of *L. lupina* using restriction enzymes. They consistently found heterozygosity in the fruiting bodies but not in the thalli, which is indicative of outcrossing and concordant with a dominant haploid life stage. Moreover, we detected very few specimens in our regional sampling with both mating types by PCR. Hence, it appears that the majority of *Letharia* thalli in nature are in fact haploid.

The conspicuous pattern of divergent haplotypes within both our and the McKenzie *L. lupina* metagenomes is consistent with hybridization. A triploid-like hybrid could be formed in a number of ways, but here we highlight some possible models (fig. 9). It has been proposed before that multiple genotypes could coexist within a single lichen thallus, forming chimeras (Jahns and Ott 1997; Dyer et al. 2001; Mansournia et al. 2012), which in the case of the triploid-like *L. lupina* thalli would be of three individuals of two divergent lineages (fig. 9Ai). However, the read count ratios along the genome are perfectly aligned to triploidy in all three, individually prepared metagenome samples, and there is no reason to expect that different genotypes growing together in a single thallus without cell fusion would maintain such strict proportions of hyphal biomass and DNA content in different thalli. Rather, it is more plausible that the metagenomes are indeed triploid-like in one of three nuclear configurations: 3n, 2n + n, or n+n+n. In plants and animals, hybridization is a common cause of triploidization via fertilization by unreduced gametes, which leads to triploid offspring that are often unstable and aneuploid (Köhler et al. 2099; Mable 2003). However, the absence of diploid-like variants in our data suggest that there is no obvious aneuploidy, and hence the triploid-like configuration is genome-wide. Moreover, the dominant life stage of fungi is haploidy, and hence the products of meiosis are directly the offspring (spores). It follows that triploidization might occur during meiosis and spore development.

**Figure 9.**
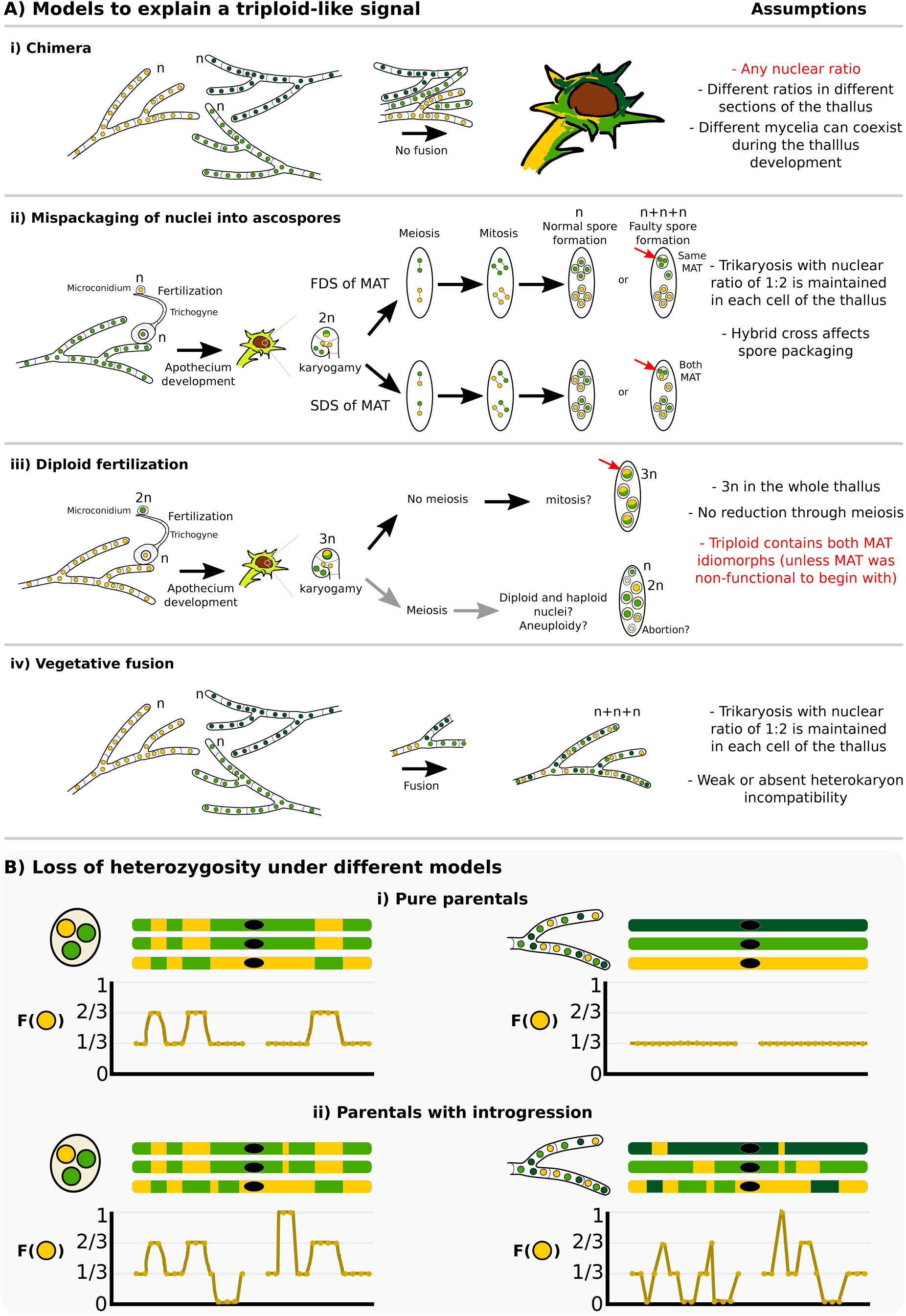
How is a triploid-like hybrid of *Letharia* formed? (**A**) We present four main models that can explain a triploid-like pattern where only two out of three (sub)genomes belong to the same species. More complex models or combinations are also possible. The *L. lupina* allele ancestry is represented by green and the unknown *Letharia* lineage by yellow. Two shades of green, light and dark, correspond to different variants of the same species. In model (**i**) chimeric thalli formed from three independent haploid (n) mycelia that do not undergo cell fusion. In model (**ii**) normal sexual reproduction with fertilization involves haploid parents, followed by a standard meiosis but with mispackaging of nuclei during ascospore formation, creating trikaryotic (n+n+n) spores. These spores can have a single mating type if the MAT locus undergoes first division segregation (FDS), or both mating types if it follows second division segregation (SDS). Model (**iii**) implies sexual reproduction where one of the parents is diploid (2n), leading to a triploid (3n) zygote during karyogamy, and a failure of meiosis to preserve the triploid condition in the resulting spores. In the last model (**iv**), vegetative fusion of three individuals leads to trikaryotic mycelia. For each model, a number of testable assumptions is given, highlighting in red those that are unlikely or do not conform to the observed data. Model (**i**) is unlikely given the triploid-like observation in three independently sequenced lichen thalli (ours and McKenzie’s). Model (**iii**) requires multiple events (e.g. previous diploidization of one of the parentals, no meiotic reduction, a bypass of the heterothallic system to produce spores of the same mating type in case of the McKenzie *L. lupina*), and given the occurrence of meiosis after karyogamy, aneuploidy is likely to occur in the resulting spores, which is also incompatible with the observed data. Hence, we interpret the models (**ii**) and (**iv**) as the most likely hypotheses, since they can accommodate thalli with a single and two MAT idiomorphs. Still, even under those two models, previous introgression between the two parentals is necessary in order to recover the observed areas of loss of heterozygosity along the genome, as shown by schematic representations in (**B**). The frequency of the yellow allele is shown along cartoon chromosomes in the absence (**i**) or presence (**ii**) of introgression. The black dot on the chromosomes represents the centromere or any repetitive area that leads to missing data.

An alternative explanation, to our knowledge absent from the literature, is the faulty packaging of nuclei into ascospores (fig. 9Aii). Normally, the packaging of nuclei during spore development is strictly controlled, but it is known from model ascomycetes like *Podospora* and *Neurospora* that occasionally this fails, and spores with an unexpected number of nuclei are accidentally formed at low rates (Ames 1934; Page 1936; Raju and Newmeyer 1977; Raju 1992). Crosses between different species have been shown to induce or exacerbate irregularities in spore development (Perkins et al. 1976; Perkins 1994; Jacobson 1995). Notably, our *L. lupina* and the McKenzie *L. columbiana* metagenomes contain both mating types (in a 1:2 ratio), but the McKenzie *L. lupina* metagenome has only one. These two types of triploid-like cases could be formed by differential segregation of the MAT locus along the ascus: under first division segregation, all nuclei in an accidentally trikaryotic spore (two mitotic sister nuclei plus one non-sister nucleus) can be of the same mating type, while under second division segregation such a spore would have two sister nuclei of one mating type and a non-sister nucleus of the other mating type (fig. 9Aii). Yet another model where fertilization occurs from one diploid parent would require the absence of meiosis to preserve the triploid (3n) configuration in the spores (fig. 9Aiii). However, given a functional heterothallic life-style, triploid spores should always have both mating types, which does not match the observation in the McKenzie *L. lupina* metagenome. Finally, the triploid-like lichens could be produced via vegetative fusion of different individuals of the ascomycete lichen symbiont (fig. 9Aiv), forming heterokaryons (Tripp and Lendemer 2018). Nonetheless, fusion of ascomycetes not involved in lichens typically leads to dikaryons (n+n) only. In the absence of knowledge on vegetative incompatibility systems in lichenized fungi, it is unclear how likely heterokaryosis would lead to trikaryons (n+n+n).

Note that these different models lead to specific predictions on the distribution of allele frequency variation along the genome (fig. 9B). For example, in the trikaryotic spore two of the genotypes should be identical, as they are the sister products of mitosis, and there should be perfectly aligned tracks of swapped ancestry between the subgenomes produced by recombination (left side of fig. 9B). This would not be the case in the heterokaryon produced by fusion unless the individuals involved are siblings (right side of fig. 9B). Moreover, the trikaryotic spore, the heterokaryon produced by fusion, and even the chimera would not be able to recover the observed distribution of allele frequencies in the absence of introgressed tracks in the parentals (compare fig.9Bi with fig. 9Bii). With the available data, it is not possible to distinguish between these different scenarios, but our results strongly suggest that there is extensive hybridization and introgression between *L. lupina* and an additional *Letharia* lineage. In addition, it remains unclear if triploid-like *Letharia* individuals are infertile or if the presence of both mating types allows for self-compatibility.

Historically, the taxonomy of the *Letharia* genus has been chronically plagued by cryptic lineages and blurry species definitions (Kroken and Taylor 2001a; Altermann 2009; Altermann et al. 2014; Altermann et al. 2016), partially due to their recent diversification (judging by their high genetic similarity), which inevitably leads to high incomplete lineage sorting. It is likely that hybrids and gene flow further complicate matters. Clearly, *L. lupina* is not the only taxon producing triploid-like thalli. Specifically, while we found that our *L. columbiana* metagenome is haploid, the sample of McKenzie et al. (2020) turned out to be triploid-like. In addition, one of the *L. vulpina* samples assayed here with PCR had both mating types. Hence, it is possible that hybridization and formation of triploid-like individuals occurs in other lineages of the genus, contributing further to the taxonomical difficulty in this group.

In a large-scale study, Pizarro et al. (2019) found that a single MAT idiomorph was found in representative samples across Lecanoromycetes, consistent with widespread heterothallism. As expected, all *Letharia* lineages studied here also follow this pattern. We further characterized both idiomorphs and determined the size of the non-recombining region (3.8 kbp), which has not been studied but for a handful of taxa (Scherrer et al. 2005; Dal Grande et al. 2018; Armaleo et al. 2019; Pizarro et al. 2019). Moreover, we refined the annotation of the auxiliary genes present in *Letharia* using transcriptomic data and homology searches to other fungal classes. This analysis confirmed that synteny is conserved in both idiomorphs for Parmeliaceae, Cladoniaceae, and Umbilicariaceae, and potentially for other families within Lecanoromycetes, making *Letharia* an adequate reference for the mating type architecture of this group. However, the MAT locus in Collemataceae and Icmadophilaceae seems to be different, highlighting that the MAT locus architecture is not conserved across all lichen-forming Lecanoromycetes.

Since heterothallism implies sexual self-incompatibility, the relative proportion of mating-types in a population is critical for a population’s ability to reproduce sexually (Consolo et al. 2005; Olarte et al. 2012; Saleh et al. 2012; Singh et al. 2012). Here we determined that both sorediate species, *L. lupina* and *L. vulpina*, likely reproduce sexually often enough to maintain balanced MAT frequencies in North American populations, despite the fact that apotheciate thalli are relatively rare. In Europe, nonetheless, the MAT frequencies are biased, to the point that only one mating type was found in the Swedish *L. vulpina* populations. The extreme scarcity of apothecia and low genetic variation in Sweden (Högberg et al. 2002) could be related to the lack of mating partner, as the populations may be pushed to predominantly clonal reproduction. Alternatively, the absence of the MAT1-1 idiomorph might reflect the preferred route of reproduction as mainly clonal propagation can skew the local distribution of mating types even if the populations rarely recombine (Singh et al. 2015). Clonality at the edge of a species’ geographical range is commonly observed across different organism groups and can have ecological advantage over short time (e.g., Billingham et al. 2003; Kawecki 2008; Meloni et al. 2013). In the Swedish range edge populations of *L. vulpina*, clonality may be especially beneficial as it preserves the combination of the compatible symbionts adapted to the local habitat, in addition to reassuring the survival of the individual and the species. However, despite the short-term advantages of clonality, the lack of recombination should eventually lead to limited evolutionary potential of the species and accumulation of deleterious mutations (Kawecki 2008). Currently, there seems to be no, or very low, potential for sexual reproduction of *L. vulpina* in the studied Swedish populations, which in combination with the low genetic diversity may reduce the fitness and the adaptive potential of these lichens in a changing environment.

## Conclusions

Hybridization and polyploidization are emerging as significant influencers of fungal evolution and speciation, based largely on the study of pathogens or human-associated species (Allman et al. 2011; Marcet-Houben and Gabaldón 2015; Depotter et al. 2016; Steenwyk et al. 2020). The results presented here, along with recent studies (Keuler et al. 2020; McKenzie et al. 2020), suggest that lichen fungal symbionts are also subject to these phenomena. The genus *Letharia* is a classical model of evolutionary biology amongst lichen-forming species thanks to the work of S. Kroken, J. W. Taylor and others (Kroken and Taylor 2001a; Kroken and Taylor 2001b; Högberg et al. 2002; Altermann et al. 2014; Altermann et al. 2016). Now, *Letharia* is elevated into the genomics era, refueling research on lichen reproductive biology. In the process new aspects of its biology are discovered, which are indeed enigmatic.

## Supplementary Figure and Table Captions

**Supplementary Figure 1.**
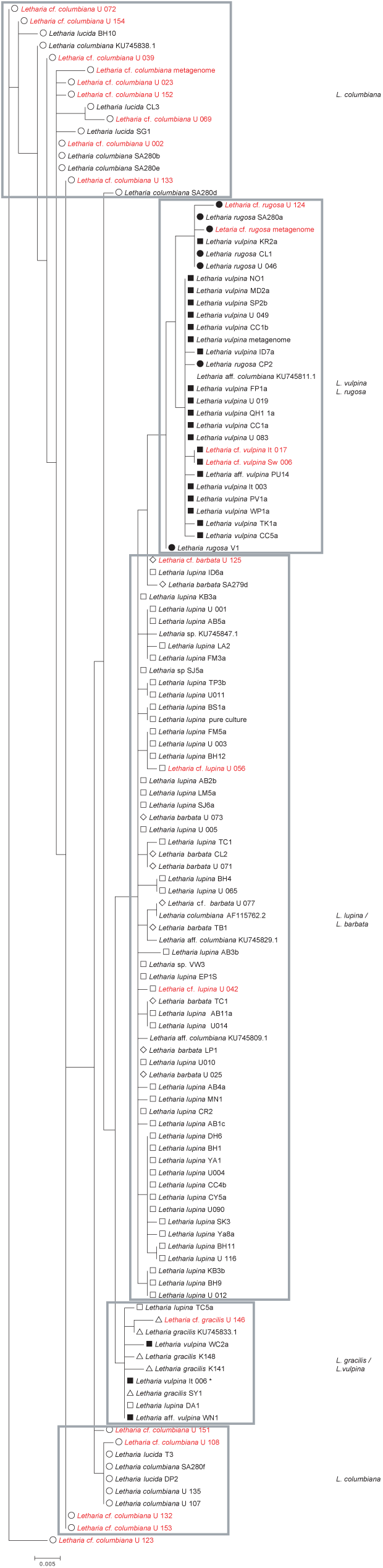
Unrooted maximum likelihood tree of ITS with all obtained variants from this study (marked with U, Sw or It) and previously published ones, the code referring to the sequences from Kroken & Taylor (2001a), Alterman et al. (2014), Alterman et al. (2016) or NCBI accession number. The unique variants obtained in this study are marked red. The Italian variant marked with * is assigned to *L. vulpina* based on algal ITS. Branch lengths are proportional to the scale bar (nucleotide substitutions per site).

**Supplementary Figure 2.**
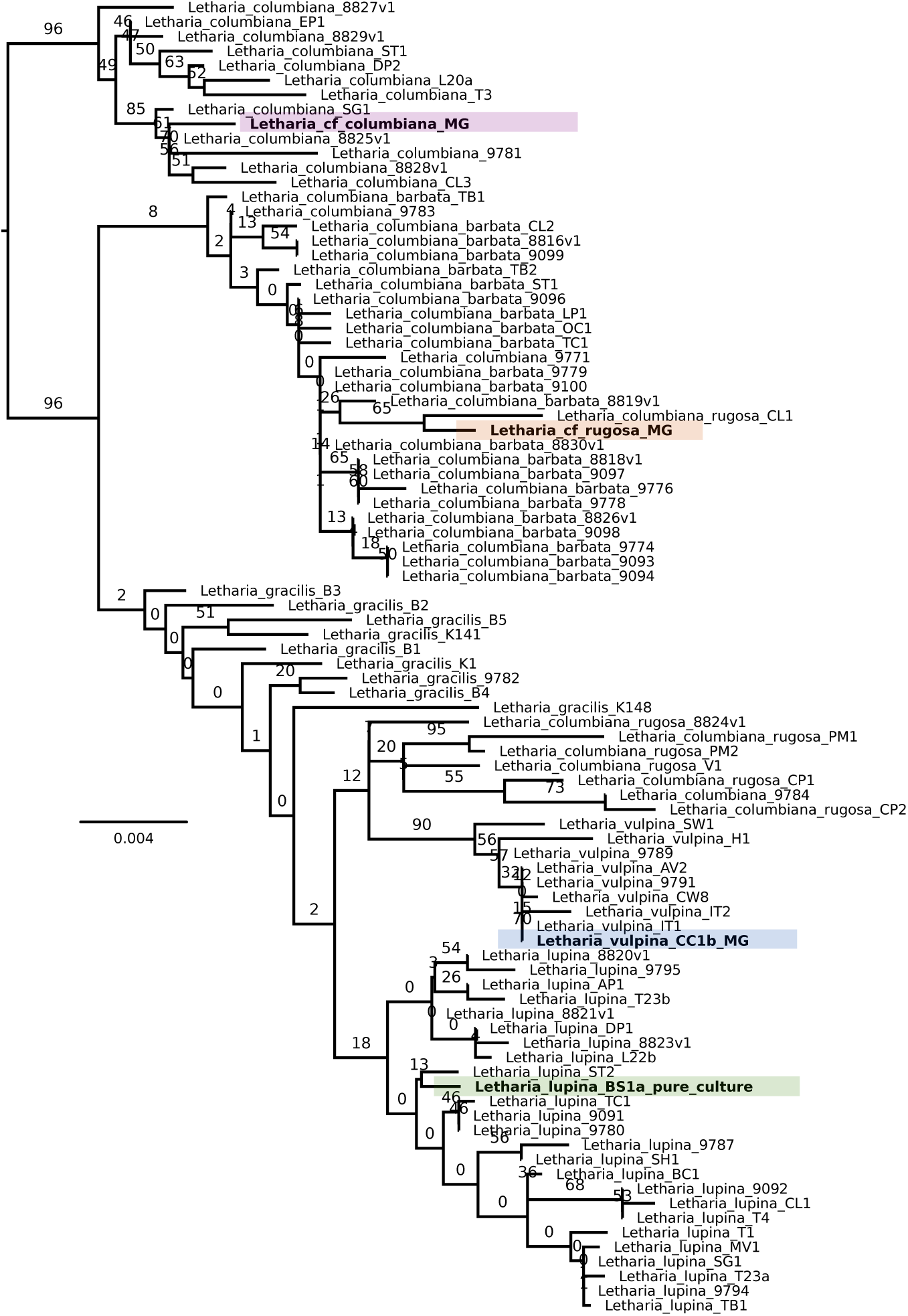
Maximum likelihood phylogeny of the *Letharia* genus produced from 11 concatenated markers. Colours highlight specimens with our Illumina data and other sequences are from previous studies. Numbers above branches represent bootstrap support values. Branch lengths are proportional to the scale bar (nucleotide substitutions per site). The tree was rooted arbitrarily with the highly supported clade containing the metagenome of *L. columbiana*. MG: metagenome.

**Supplementary Figure 3.**
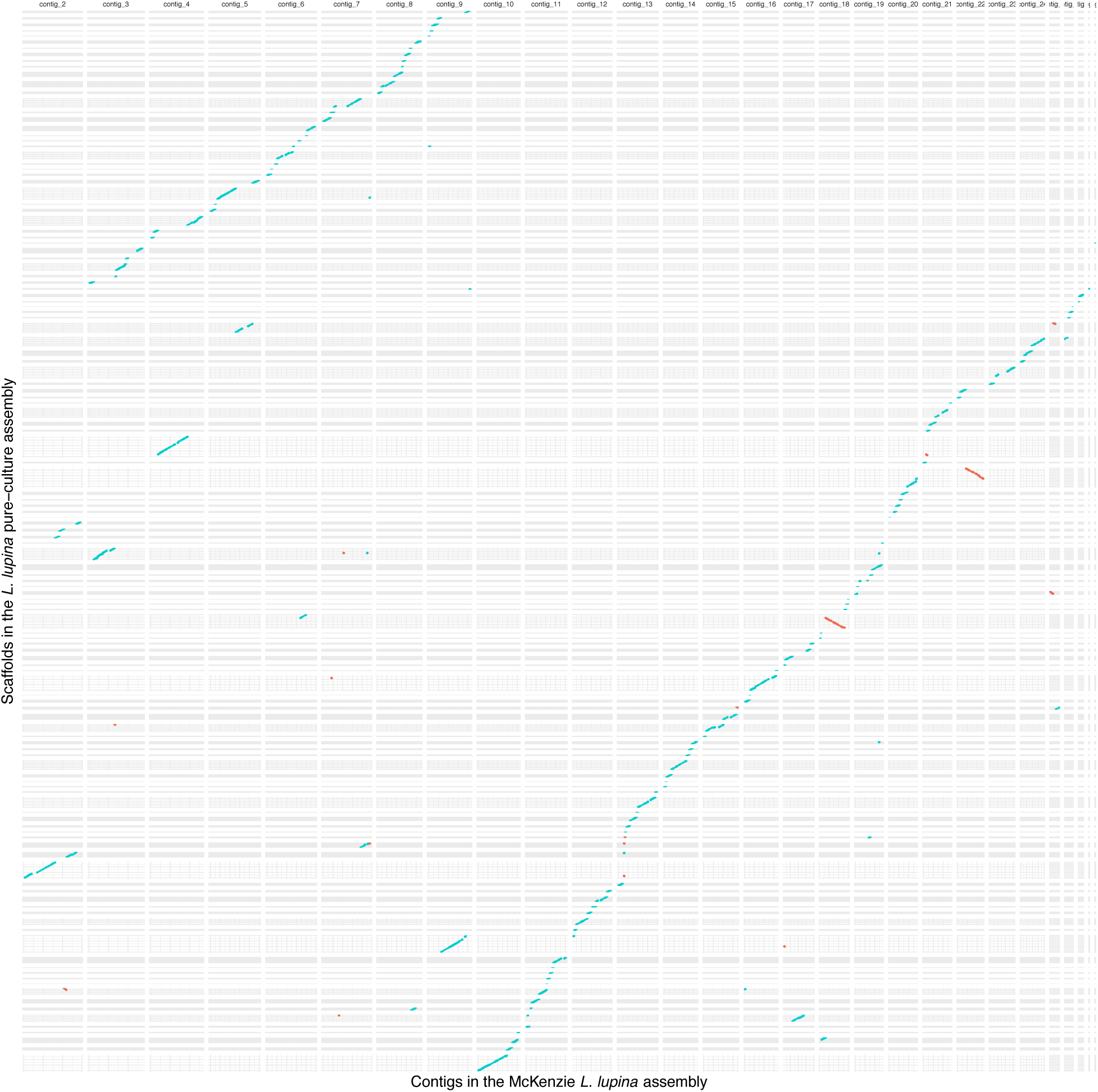
Dotplot of MUMmer alignments between the assemblies of the McKenzie *L. lupina* metagenome and our *L. lupina* pure culture. Scaffold names of the pure culture assembly are omitted for clarity. Blue indicates alignments in the same sense, while red marks inversions. The Contig 1 of the McKenzie *L. lupina* assembly is excluded since alignments are too short and scarce to be visible.

**Supplementary Figure 4.**
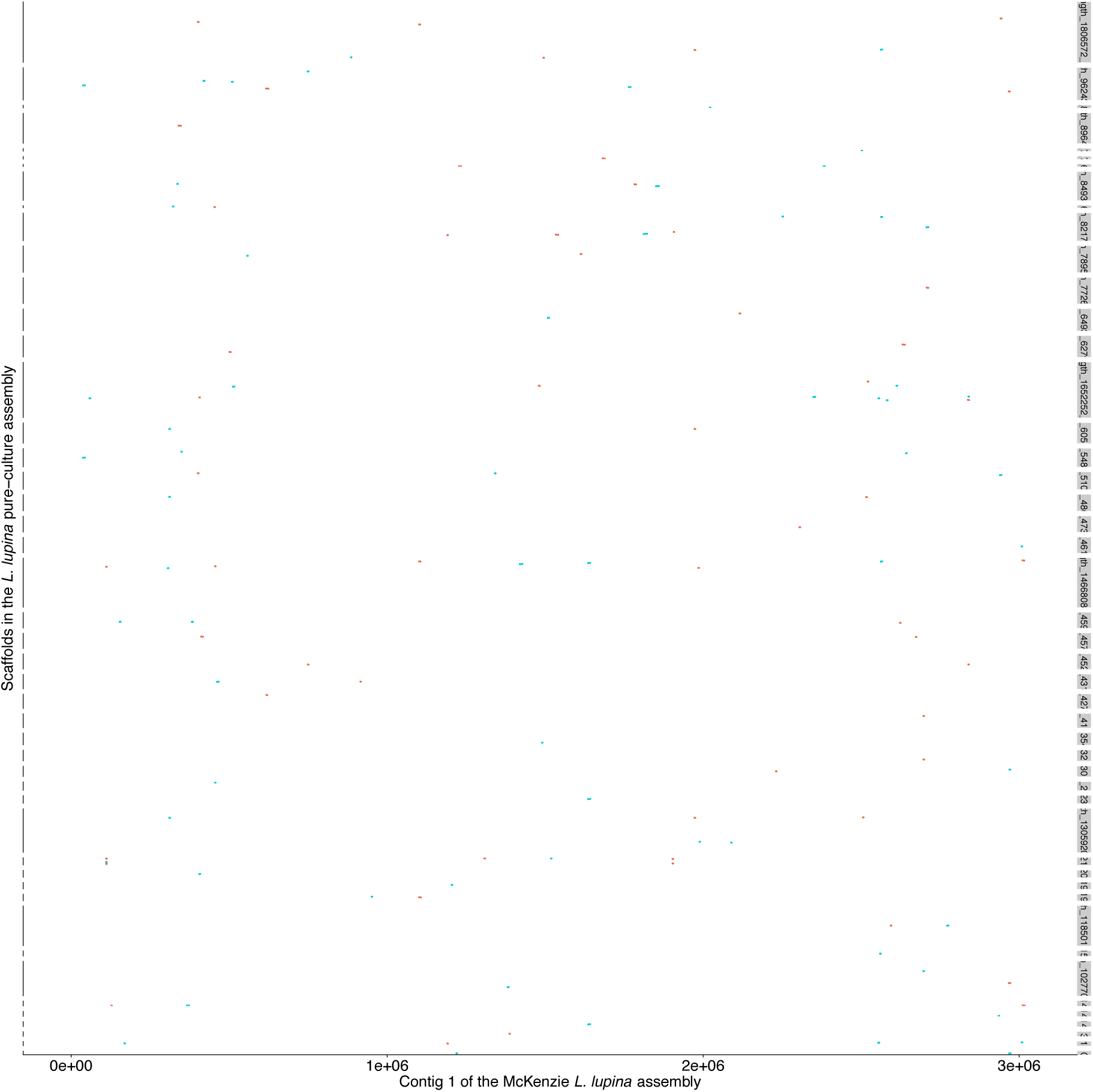
Dotplot of MUMmer alignments between the Contig 1 of the McKenzie *L. lupina* metagenome and our *L. lupina* pure-culture assembly. Blue indicates alignments in the same sense, while red marks inversions.

**Supplementary Figure 5.**
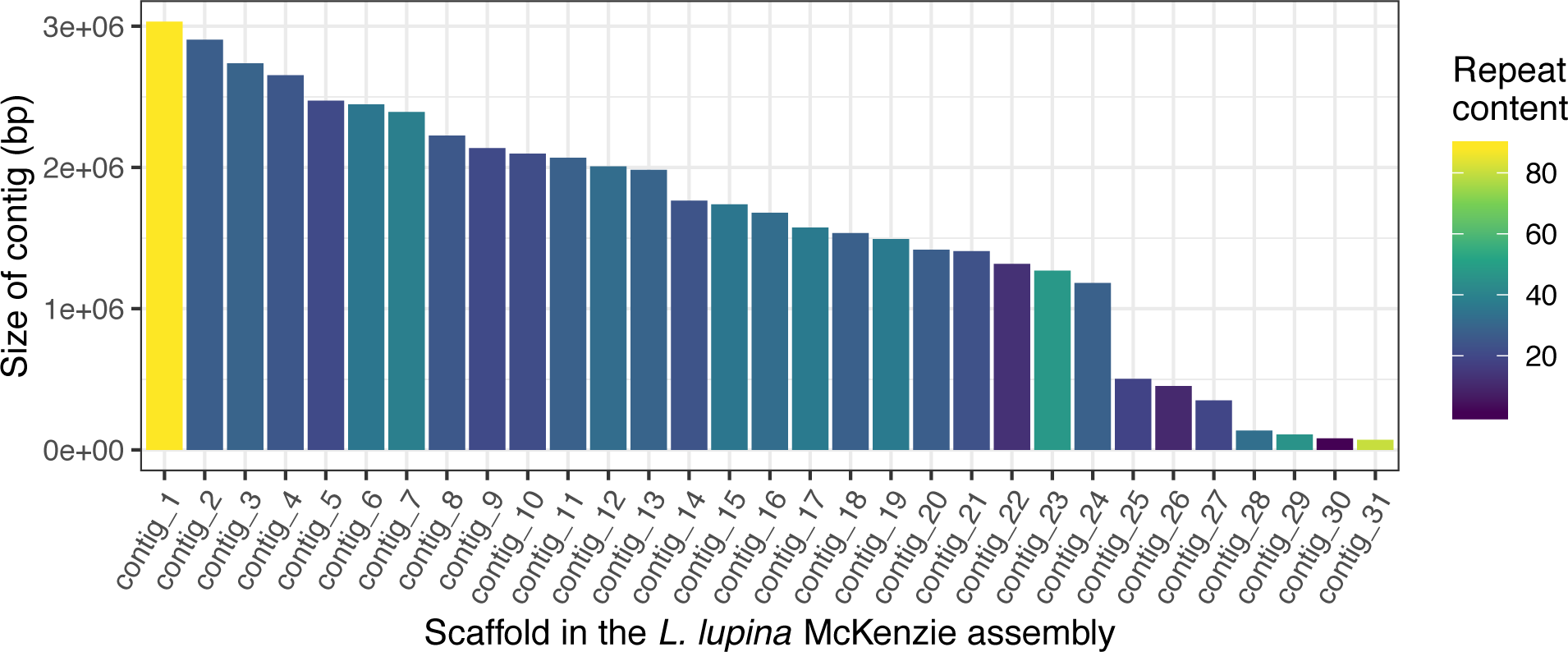
Repeat content proportions of the contigs in the McKenzie *L. lupina* metagenome assembly.

**Supplementary Figure 6.**
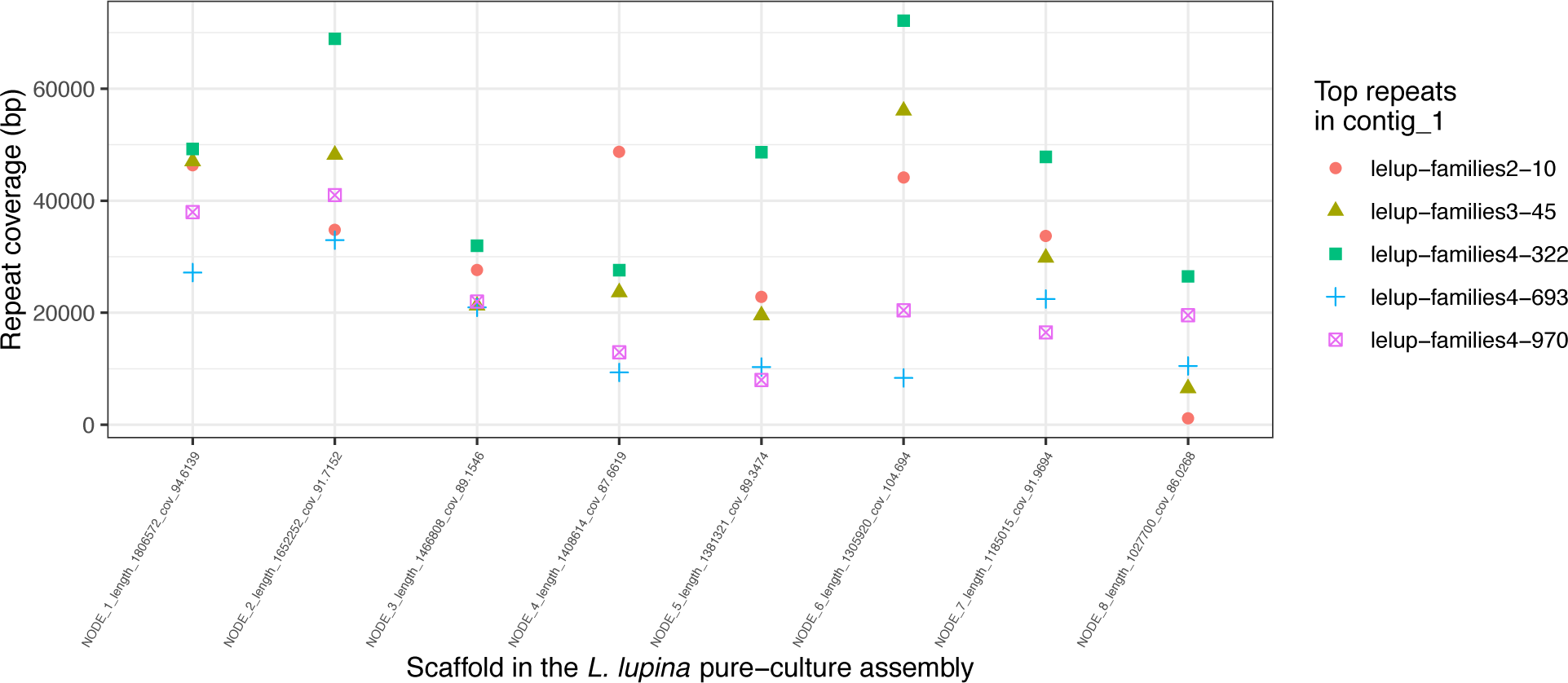
Coverage (in total number of base pairs) of the top five most abundant repetitive elements in the Contig 1 from the McKenzie *L. lupina* metagenome across the biggest scaffolds of the *L. lupina* pure culture assembly.

**Supplementary Figure 7.**
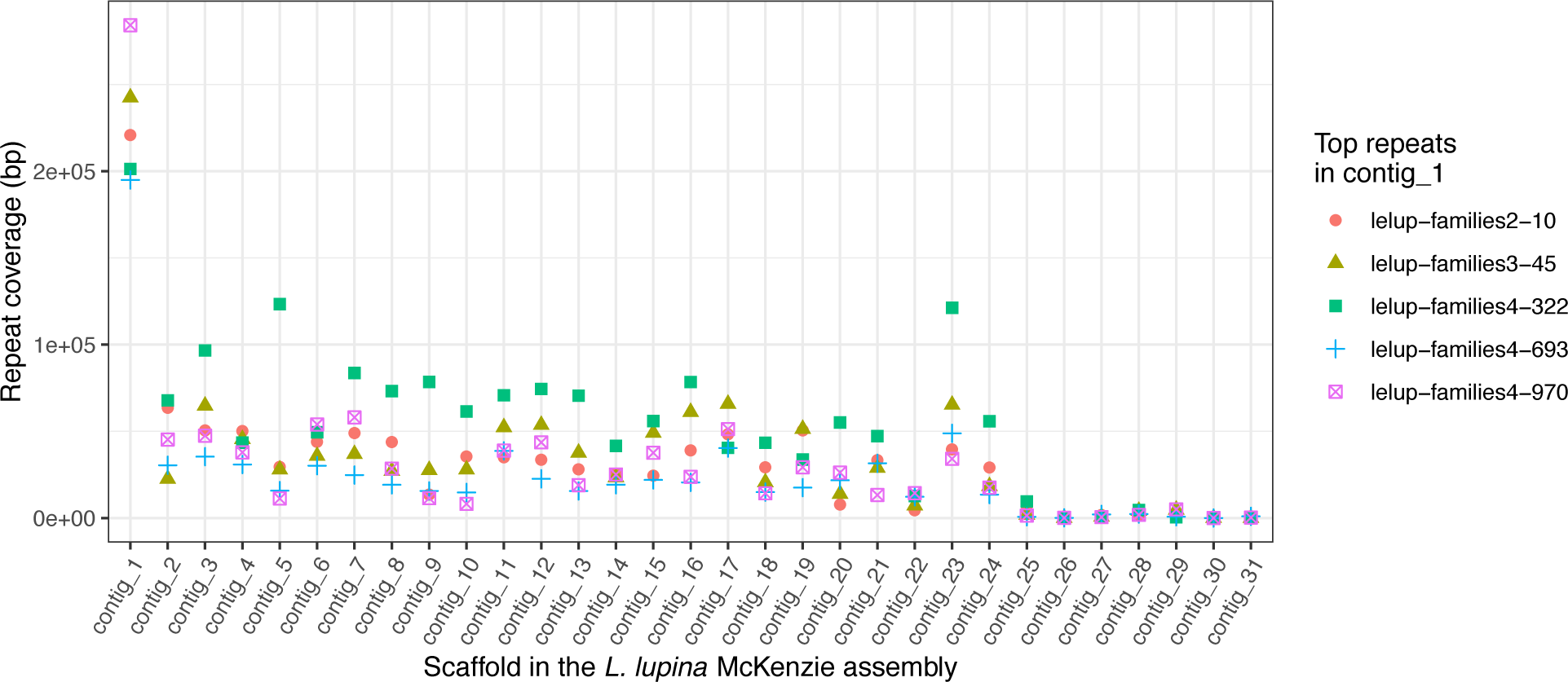
Coverage (in total number of base pairs) of the top five most abundant repetitive elements in the Contig 1 across all the other contigs of the McKenzie *L. lupina* metagenome assembly.

**Supplementary Figure 8.**
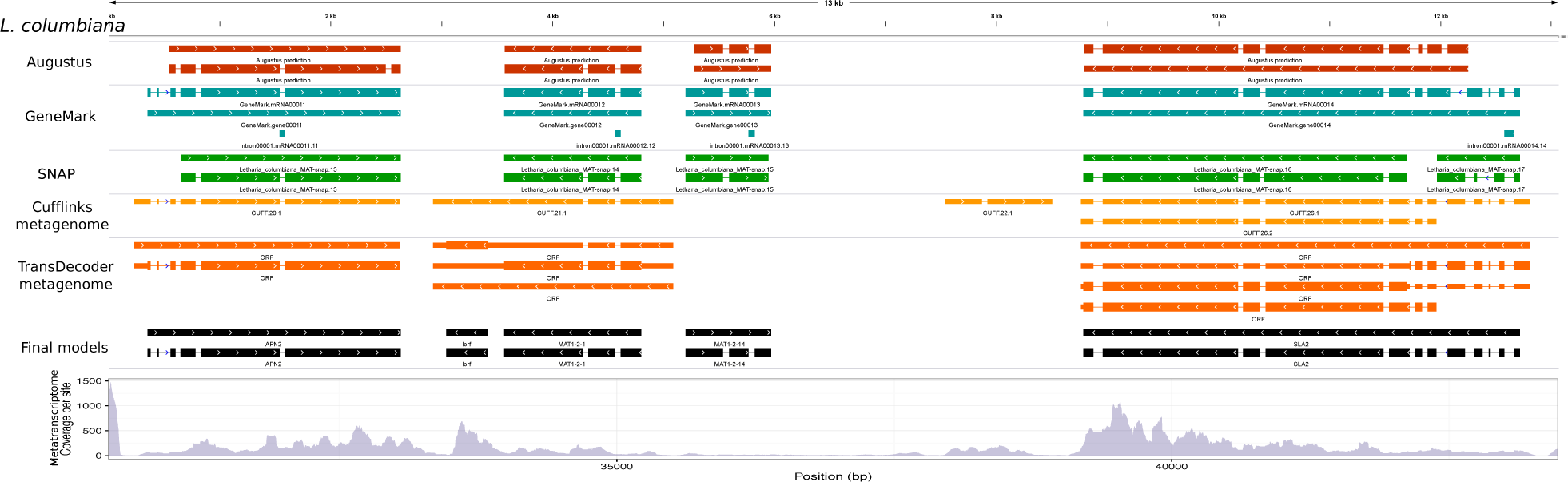
Annotation of the MAT locus of *Letharia columbiana*. The main sources of evidence for annotation are shown. These include the *ab initio* gene models (Augustus, GeneMark and SNAP), transcript models (Cufflinks), and ORFs from transcripts (TransDecoder). The final models are presented in black. At the bottom, we present the depth of coverage per site from the metatranscriptome.

**Supplementary Figure 9.**
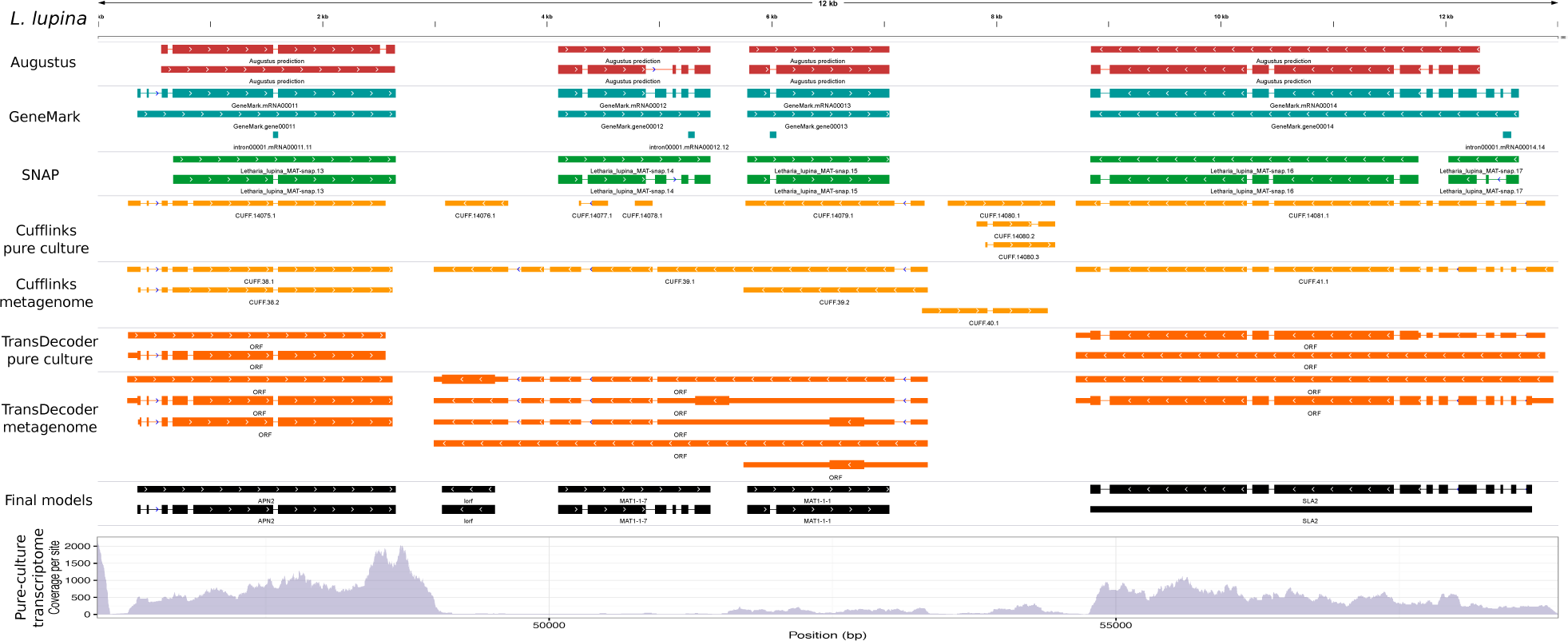
Annotation of the MAT locus of *Letharia lupina*. The main sources of evidence for annotation are shown. These include the *ab initio* gene models (Augustus, GeneMark and SNAP), transcript models (Cufflinks), and ORFs from transcripts (TransDecoder). The final models are presented in black. Note that the transcriptomic models and ORFs for both the pure culture and the lichen thallus (metagenome) are shown. The depth of coverage at the bottom is only for the pure culture.

**Supplementary Figure 10.**
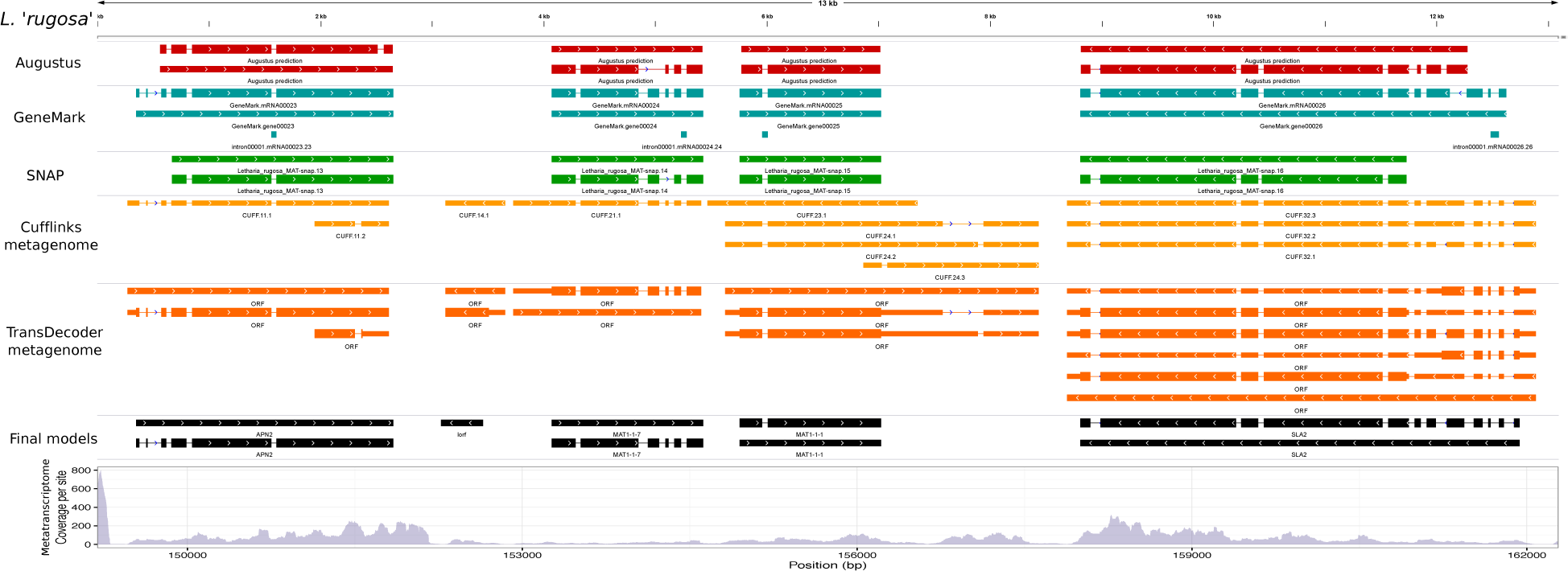
Annotation of the MAT locus of *Letharia ‘rugosa’*. The main sources of evidence for annotation are shown. These include the *ab initio* gene models (Augustus, GeneMark and SNAP), transcript models (Cufflinks), and ORFs from transcripts (TransDecoder). The final models are presented in black. At the bottom, we present the depth of coverage per site from the metatranscriptome.

**Supplementary Figure 11.**
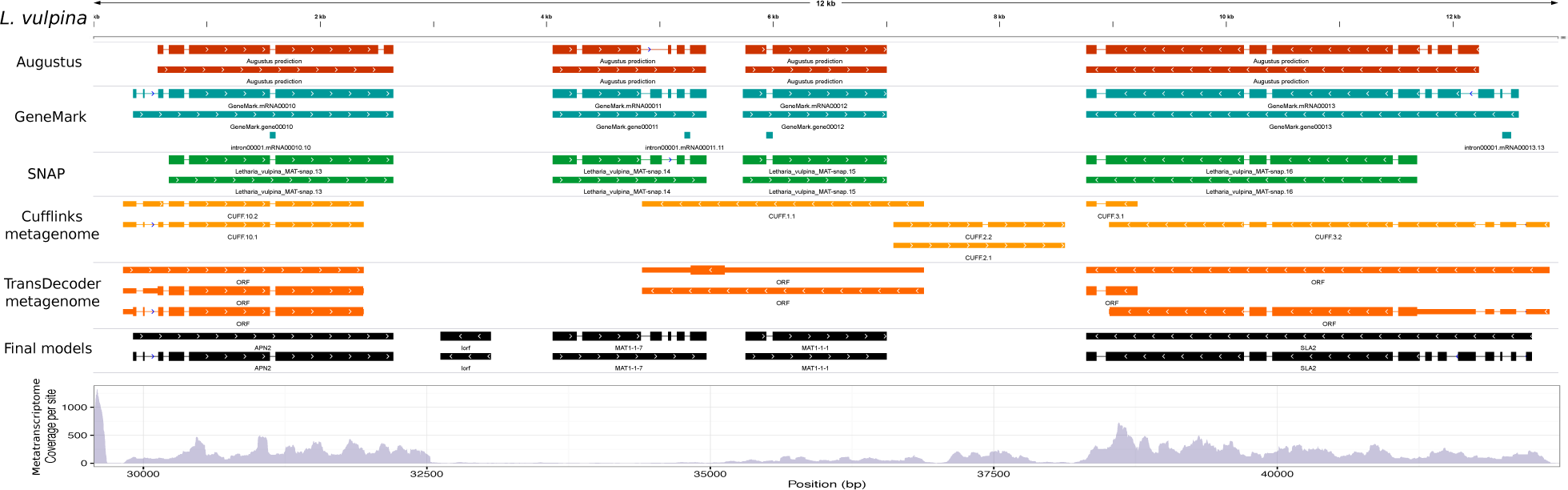
Annotation of the MAT locus of *Letharia vulpina*. The main sources of evidence for annotation are shown. These include the *ab initio* gene models (Augustus, GeneMark and SNAP), transcript models (Cufflinks), and ORFs from transcripts (TransDecoder). The final models are presented in black. At the bottom, we present the depth of coverage per site from the metatranscriptome.

**Supplementary Figure 12.**
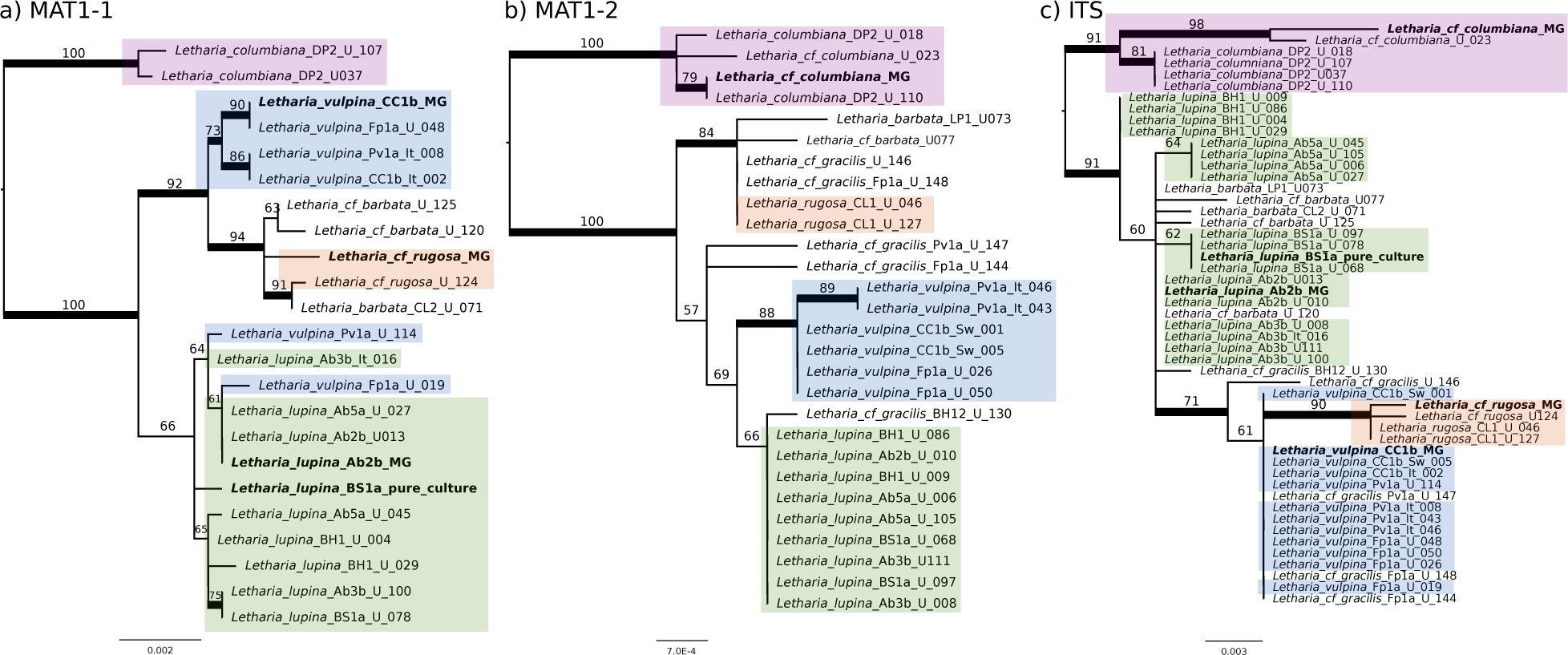
(previous page) Maximum likelihood trees of the alignment of (**A**) MAT1-1 (encompassing the *MAT1-1-7* and *MAT1-1-1* genes along with their intergenic region), MAT1-2 (*MAT1-2-1* and *MAT1-2-14* with their intergenic region), and the fungal barcode ITS (**C**). Bootstrap values are shown above the branches. Branches with bootstrap values above 70% are thickened. Colours highlight the four main taxa under study: *L. columbiana* in purple, *L. ‘rugosa’* in orange, *L. vulpina* in blue and *L. lupina* in green. Samples with genomic or metagenomic data are in bold. Trees were rooted arbitrarily with *L. columbiana.* Branch lengths are proportional to the scale bar (nucleotide substitutions per site).

**Supplementary Figure 13.**
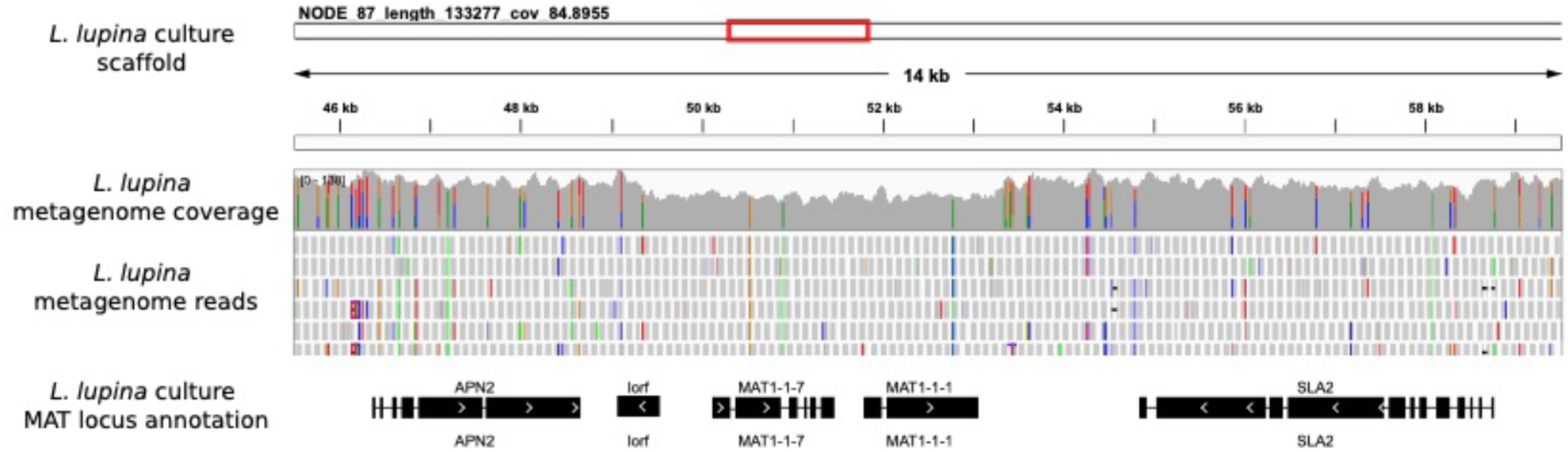
Depth of coverage of the *L. lupina* metagenome reads mapped along the MAT idiomorph as displayed by the Integrative Genomics Viewer (IGV) program. The pure culture assembly (MAT1-1) was used as reference. The coverage within the idiomorph drops to approximately two thirds of the normal coverage, as expected for a 2:1 proportion of the MAT1-1 and MAT1-2 subgenomes within the *L. lupina* metagenome data. Colored columns along the coverage track represent the polymorphic sites and their frequencies.

**Table S1**. *Letharia* specimens used in this study and detailed information on their collection sites, obtained MAT idiomorphs and data repository. The taxon identification is based on the ITS variants and thallus morphology. The ITS variants are named with the codes used by Kroken and Taylor (2001a) and Altermann et al. (2014).

**Table S2.**
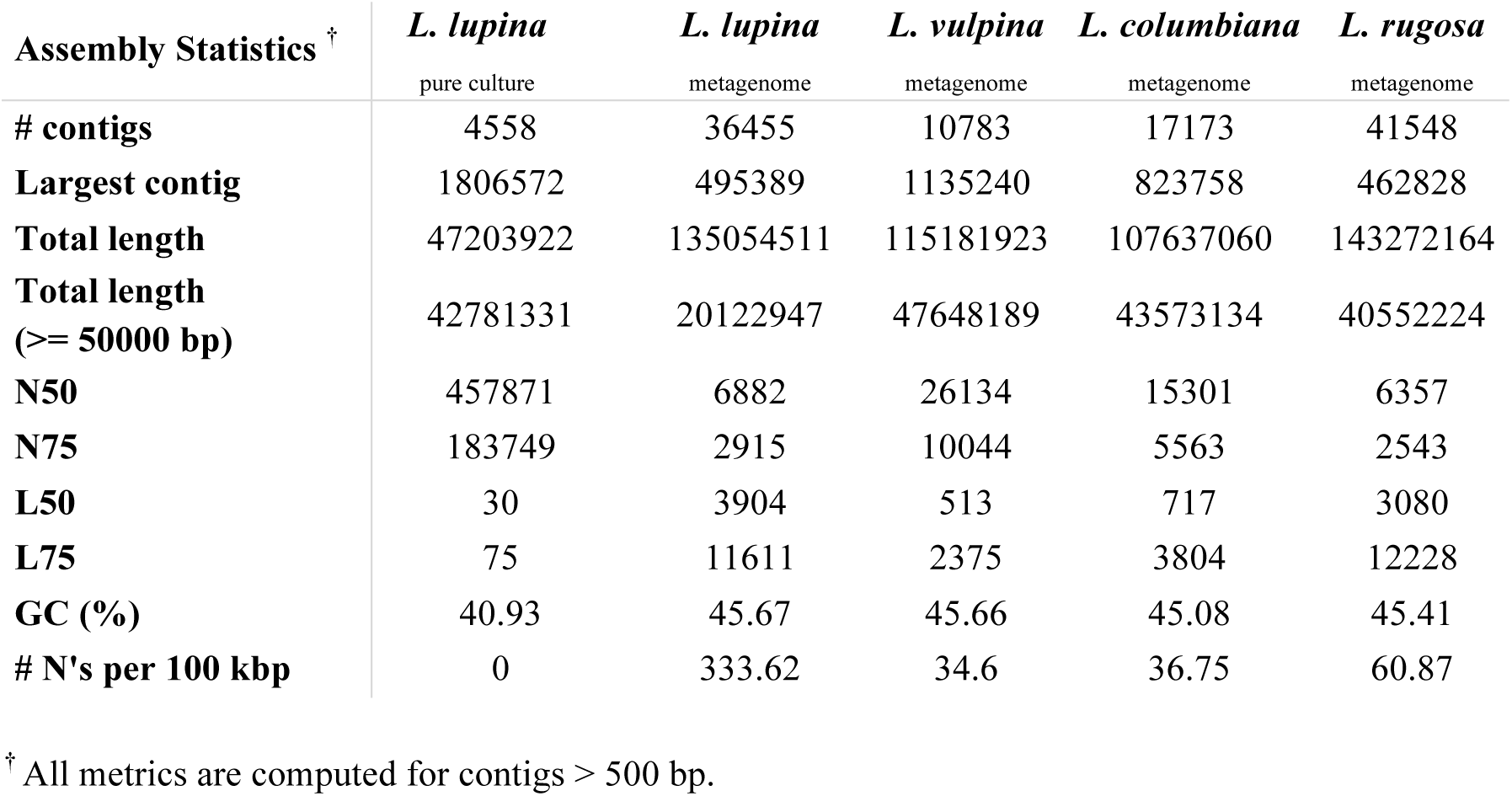
Assembly statistics of the draft assemblies used in this study.

| Assembly Statistics <sup>†</sup> | *L. lupina* | *L. lupina* | *L. vulpina* | *L. columbiana* | *L. rugosa* |
| --- | --- | --- | --- | --- | --- |
|  | pure culture | metagenome | metagenome | metagenome | metagenome |
| **# contigs** | 4558 | 36455 | 10783 | 17173 | 41548 |
| **Largest contig** | 1806572 | 495389 | 1135240 | 823758 | 462828 |
| **Total length** | 47203922 | 135054511 | 115181923 | 107637060 | 143272164 |
| **Total length**<br>**(≥ 50000 bp)** | 42781331 | 20122947 | 47648189 | 43573134 | 40552224 |
| **N50** | 457871 | 6882 | 26134 | 15301 | 6357 |
| **N75** | 183749 | 2915 | 10044 | 5563 | 2543 |
| **L50** | 30 | 3904 | 513 | 717 | 3080 |
| **L75** | 75 | 11611 | 2375 | 3804 | 12228 |
| **GC (%)** | 40.93 | 45.67 | 45.66 | 45.08 | 45.41 |
| **# N's per 100 kbp** | 0 | 333.62 | 34.6 | 36.75 | 60.87 |

<sup>†</sup> All metrics are computed for contigs > 500 bp.

**Table S3.**
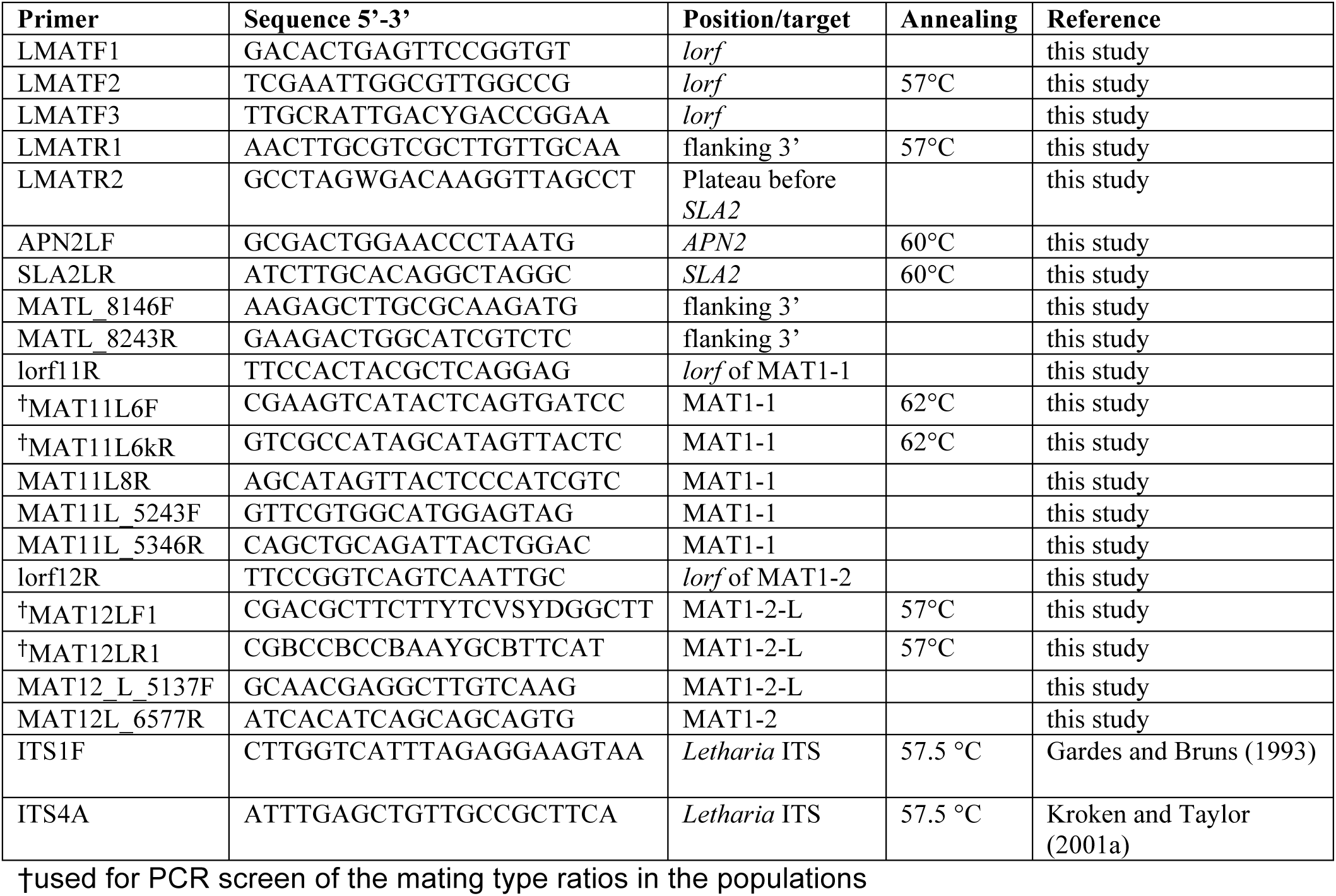
Primers used for the PCR amplification and sequencing of the MAT locus in *Letharia*. The primers with no annealing temperature were used only for sequencing. The ITS primers were used to amplify the ITS region for all studied specimens (Tuovinen et al., 2019).

## Acknowledgments

This project was supported by a stipend to LB by Leksand municipality, grants from Stiftelsen Lars Hiertas minne and Stiftelsen Extensus to VT, and by a grant (DO2011-0022) from Stiftelsen Oscar och Lili Lamms minne to GT. We acknowledge that the sequencing of metagenomes and pure culture was performed by the SNP&SEQ Technology Platform, Science for Life Laboratory at Uppsala University, a national infrastructure supported by the Swedish Research Council (VR-RFI) and the Knut and Alice Wallenberg Foundation. A collecting permit for *Letharia vulpina* in Sweden was issued by the county administration board of Dalarna (Dnr: 522-2513-2014). Helmut Mayrhofer and John McCutcheon are thanked for supporting metatranscriptome sequencing at University of Graz and University of Montana, respectively. We thank Bruce McCune and Steve Leavitt for providing us with *Letharia* specimens, Markus Hiltunen for laboratory assistance, and Gulnara Tagirdzhanova, Sergio Tusso, and Aaron A. Vogan for fruitful discussions about the metagenomes and analyses, and Dr. Markus Wilken about the MAT nomenclature. We also thank Jessica L. Allen and Sean McKenzie for access to their *Letharia* repeat library and genome annotations.

## Data accessibility

The raw genome and transcriptome data are available through NCBI’s Short Read Archive with accession PRJNA523679. The annotated MAT loci are available in GenBank with accessions MK521629-MK521632. The alignments used for the phylogenetic analyses are available in Dryad: XXXX. The custom scripts written to analyse the data are publicly available on https://github.com/johannessonlab/Letharia.

## Author contributions

VT, LB and HJ designed the study. VT and LB collected the Swedish specimens, TS the American specimens, JN the Italian specimens, HJ the Swiss specimens and YY provided the pure culture of *Letharia*. VT, LB, DV and TS conducted the laboratory work. SLAV, VT, LB, and DV conducted the data-analyses. GT and HJ provided the research resources. SLAV, VT and HJ wrote the paper. TS, DV, JN and GT commented and contributed to the final version of the manuscript.

## Appendix

### Identification of *Letharia* specimens

We used the ITS marker as a barcode, together with thallus morphology (sorediate vs. apotheciate) to assign *Letharia* thalli to the previously described species (Kroken and Taylor 2001a; Altermann 2009; Altermann et al. 2016). In addition, we found 18 unique ITS variants (**Table S1**). The specimens with unique ITS variants were putatively assigned to previously known lineages based on thallus morphology and ML analyses of the ITS and marked with “cf.” in fig. S1. The ITS alignment, including variants unique to this study, and previously published variants, was 555 bp long with 69 variable sites. The ITS trees are in concordance with previously published results (Kroken and Taylor 2001a; Altermann et al. 2014; Altermann et al. 2016). Two Italian specimens had a 100% identical match to previously published ITS variants extracted from *L. gracilis*, *L. vulpina* and *L. lupina* (Altermann 2009). These specimens were putatively assigned to *L. vulpina* based on the identity of the symbiotic alga from the thallus (ITS sequence, data not shown), following Altermann (2009) and Altermann et al. (2016). We had eight specimens putatively assigned to *L. gracilis* in our dataset (morphology-based identification), but the ITS variants for these did not match the previously published *L. gracilis* variants but instead those of *L. lupina* or *L. vulpina* (**Table S1**). These specimens were designated *L. cf. gracilis* and their species identity cannot be resolved with current data.

### Training of *ab initio* gene predictors for *Letharia* genomes

The program SNAP was trained a priori with the *L. lupina* transcripts produced by Trinity, following the instructions in the SNAP’s README and Campbell et al. (2014). The MAKER pipeline v. 2.31.8 (Holt and Yandell 2011) was run to infer gene predictions directly from the Trinity transcripts (*est2genome=1*). The resulting GFF files were transformed into the non-standard format ZFF using the script *maker2zff*. All models with warnings or errors reported by the program fathom where excluded. The program forge was used to estimate parameters for the script hmm-assembler.pl. The resulting HMM training files were used to run MAKER once more, this time with *est2genome=0*. SNAP was retrained as above using this new MAKER annotation. The HMM training files of the retraining process were used for annotation in downstream analyses. Augustus was trained using the protein sequences of the gene models produced by running SNAP on the SPAdes assembly of *L. lupina*. We used the script *autoAug.pl* (part of Augustus distribution) to do the training with *--maxIntronLen=1000*. Finally, GeneMark was self-trained using the script *gmes_petap.pl* and the flags: -*-fungal --max_intron 3000 --min_gene_prediction 120*. All scripts used and the training files are available at https://github.com/johannessonlab/Letharia.

### Additional results of the MAT region annotation

The metatranscriptomic data supported the presence of a small ORF (herein referred to as *lorf*, from *<u>L</u>etharia* <u>o</u>pen reading frame) between *APN2* and *MAT1-2-1* in *L. columbiana*, flanking the beginning of the idiomorphic region (fig. 2 and fig. S8). In the other species, however, *lorf* displays relatively low levels of expression (see fig. S8-S11), which may be due to differences in biological conditions during the time of collection. In the *L. lupina* metatranscriptome, TransDecoder detected *lorf* within the transcripts, suggesting activity in this species.

Nevertheless, an alignment of all samples (including sequences acquired from PCR) revealed a potentially on-going process of pseudogenization in the entire genus. All the *lorf* sequences present in samples of the MAT1-1 idiomorph have two potential start codons (AUG), while some *lorf* sequences next to a MAT1-2 idiomorph have either one start codon, or none. Some samples also have frame-shift mutations. Noteworthy, *lorf* is located at the border between the idiomorph and the flanking region (fig. 2) and shows a slight divergence (∼96% of similarity) between MAT1-2 (*L. columbiana*) and MAT1-1 (*L. lupina*, *L. vulpina*, and *L. ‘rugosa’*).

In addition, the *Letharia* transcript models for the *MAT1-1-1* gene were consistently predicted in the opposite sense, based on homology with other fungal species and the *ab initio* gene predictors. This antisense transcript (fig. 2A) was present in all *Letharia* species, including the transcriptome of the pure culture of *L. lupina*. We recovered the transcript in the canonical sense for *MAT1-1-1* along with antisense transcripts only for *L. ‘rugosa’* (fig. 2; see also fig. S8-S11).

### Nomenclature of auxiliary MAT genes

The nomenclature for mating-type genes as suggested by Turgeon and Yoder (2000) is used for most fungal groups. Under this system, whenever a new gene is discovered, with no detectable homology to any other, an incremental number is assigned to it following certain guidelines (Turgeon and Yoder 2000). Unfortunately, there is no centralized database associated with all published auxiliary MAT genes, which has led to a number of inconsistencies and overlaps with gene names (Wilken et al. 2017). For example, genes homologous to the auxiliary gene in *Letharia* have previously been named *MAT1-1-4*, *MAT1-1-7*, or *MAT1-1-9* (fig 3, left), depending on the author (Wilken et al. 2017). As discussed by Wilken et al., this gene is not related to the *MAT1-1-4* present in the Leotiomycete *Pyrenopeziza brassicae*, so this name is not appropriate. However, we disagree with Wilken et al. (2017) in giving a separate name for *MAT1-1-*7 and *MAT1-1-9* (and perhaps even the *MAT1-1-8* gene present in the Dothidiomycete *Shaeropsis sapinea*) on the basis of low pairwise identity, as the high divergence between these genes is coherent with the large phylogenetic distance between these fungi. Hence, we refer to the *Letharia* gene as *MAT1-1-7*, which is the smallest number available. Incidentally, Armaleo et al. (2019) also used this name for the ortholog in *Cladonia*.

Since the auxiliary gene in the MAT1-2 idiomorph of *Letharia* has no detected homologs outside Lecanoromycetes, a new number should be assigned. The highest number in use that we could identify is *MAT1-2-12*. However, two independent studies used the name *MAT1-2-12* for the genes in unrelated taxa: *Teratosphaeria* (Aylward et al. 2019) and *Calonectria* (Li et al. 2020). We confirmed that these genes are not homologous to each other, or to the gene in *Letharia*, using BLASTp searches. Hence, the correct name for the *Letharia* auxiliary gene is *MAT1-2-14* and the genes in *Calonectria* (Li et al. 2020) should be called *MAT1-2-13* based on the date of publication.

